# A whole-cortex probabilistic diffusion tractography connectome

**DOI:** 10.1101/2020.06.22.166041

**Authors:** Burke Q. Rosen, Eric Halgren

## Abstract

The WU-Minn Human Connectome Project (HCP) is a publicly-available dataset containing state-of-art structural, functional, and diffusion-MRI for over a thousand healthy subjects. While the planned scope of the HCP included an anatomical connectome, resting-state functional-MRI forms the bulk of the HCP’s current connectomic output. We address this by presenting a full-cortex connectome derived from probabilistic diffusion tractography and organized into the HCP-MMP1.0 atlas. Probabilistic methods and large sample sizes are preferable for whole-connectome mapping as they increase the fidelity of traced low-probability connections. We find that overall, connection strengths are lognormally distributed and decay exponentially with tract length, that connectivity reasonably matches macaque histological tracing in homologous areas, that contralateral homologs and left-lateralized language areas are hyperconnected, and that hierarchical similarity influences connectivity. We compare the diffusion-MRI connectome to existing resting-state fMRI and cortico-cortico evoked potential connectivity matrices and find that it is more similar to the latter. This work helps fulfill the promise of the HCP and will make possible comparisons between the underlying structural connectome and functional connectomes of various modalities, brain states, and clinical conditions.

**Significance Statement:** The tracts between cortical parcels can be estimated from diffusion MRI, but most studies concentrate on only the largest connections. Here we present an atlas, the largest and most detailed of its kind, showing connections among all cortical parcels. Connectivity is relatively enhanced between frontotemporal language areas and homologous contralateral locations. We find that connectivity decays with fiber tract distance more slowly than predicted by brain volume and that structural and stimulation-derived connectivity are more similar to each other than to resting-state functional MRI correlations. The connectome presented is publicly available and organized into a commonly used scheme for defining brain areas in order to enable ready comparison to other brain imaging datasets of various modalities.

## Introduction

In the 21^st^ century, advances in computation, theory, and neuroimaging have spurred a broad and intense interest in the anatomical connections and physiological correlations among human brain areas. Bivariate functional connectivity has given way to full functional connectomes, the most comprehensive of which may be the WU-Minn Human Connectome Project’s (HCP) resting-state fMRI dense connectome (Van Essen et al., 2013). The planned scope of WU-Minn HCP also included a full anatomical connectome (Van Essen and Ugurbil, 2017), and the project has collected, curated, and preprocessed diffusion imaging (dMRI) data for 1,065 subjects. However, a structural connectome has to-date not been released for these data. This report seeks to address this omission by presenting a full-cortex anatomical connectome derived from local, probabilistic tractography.

dMRI techniques detect white matter by registering the orientation biases of water molecule diffusion within myelinated axons. The majority of dMRI studies focus on differences in specific connections between treatment groups. In contrast, we seek here to present a robust, densely populated average connectivity matrix for the entire cortex using data from a large, healthy sample. Local dMRI fiber tract tracing algorithms can be broadly organized into two classes: deterministic e.g. dsi-studio (Yeh et al., 2013), and probabilistic e.g. probtrackX (Behrens et al., 2007). Deterministic tractography considers the most likely orientation at each voxel yielding the maximum likelihood tracts whereas probabilistic tractography considers the entire distribution of possible orientations, yielding a probability cloud of connections. As our goal is instead to explore all possible connections between regions, we employed local, probabilistic tractography (Behrens et al., 2007). This method has been validated against macaque retrograde tracers within-species (Donahue et al., 2016) and the dMRI protocol and equipment used for the WU-Minn HCP database were optimized in anticipation of this analysis (Sotiropoulos et al., 2013).

The physiological relevance of a connectome is maximized if its nodes form functionally distinct areas. Within the scope of cortex, this amounts to selecting a parcellation scheme. The HCP multi-modal parcellation (HCP-MMP1.0) (Glasser et al., 2016) has several advantages: it’s boundaries are both functionally and anatomically guided, it has sufficient parcels (360) to generate a rich connectome while few enough that the parcels’ extents comfortably exceed the dMRI voxel size, and mechanisms exist (Fischl et al., 2004) for it to be readily applied to individuals. Most importantly, the HCP-MMP1.0 parcellation is publicly available and widely adopted, facilitating the comparison of the generated matrices to other structural and functional connectomes.

Given the computational intensity of dMRI fiber tractography and the field’s inclination towards elucidating specific connections, it is not surprising that the number of existing publicly available dMRI datasets exceeds that of finished, readily applicable connectivity matrices. However, there do exist some prior examples. The USC Multimodal connectivity database (http://umcd.humanconnectomeproject.org), contains two dMRI tractography connectomes with standard surface-based parcellations: Hagmann (Hagmann et al., 2008) and ICBM (Mori et al., 2008), with sample-sizes of 5 and 138, respectively. A third is available at http://www.dutchconnectomelab.nl which contains 114 controls. All of these use the Desikan-Killiany atlas (Desikan et al., 2006) which consists of 68 cortical parcels and were produced with deterministic tractography. An atlas of major fiber tracts for the HCP 1200 cohort has recently released at http://brain.labsolver.org, (Yeh et al., 2018). However, this deterministic tractography connectome is spatially coarse, consisting of only 54 cortical parcels, and lacks dynamic range and statistical dispersion, as weaker connections are unrepresented, rendering the connectivity matrix nearly binary. The HCP-MMP1.0 atlas employed here has more than five times as many parcels while retaining the functional distinctness of areas. In contrast to the relatively sparse existing deterministic matrices, the probabilistic approach may better resolve weak or low probability connections leading to densely populated connectivity matrices like those found non-human primate tracing studies (Markov et al., 2014). Furthermore, the cohort studied is large and many other types of data are available for the same individuals including the NIH neuropsychological toolbox (Gershon et al., 2013), as well as fMRI and MEG data for resting-state and cognitive tasks, permitting within-cohort comparison to functional connectivity.

The following report presents a novel structural connectome of the human neocortex based on probabilistic diffusion tractography. The connectome is partially validated against retrograde tracing in macaques and the relationship between tract length and connection strength is quantified. Further validation is provided by reasonable connectivity properties between contralateral homologous parcels, within language cortex, and between parcels lying at similar levels of the cortical hierarchy. Finally, the dMRI connectome is compared to cortico-cortico evoke potential (CCEP) and resting-state fMRI derived connectivity.

## Materials & Methods

### Subjects & data sources

No new data was collected for this study, and the existing data used was gathered from publicly available databases. Individual subject’s high-resolution T1-weighted structural magnetic resonance volumes (MRI), Diffusion images (dMRI), and group average grayordinate resting-state function MRI (rs-fMRI) connectivity were gathered from the Human Connectome Project’s (HCP) WU-Minn 1200 release (Van Essen et al., 2013) at https://db.humanconnectome.org. The diffusion imaging dataset consists of 1065 individuals (575 women), aged 22-36+ years old. The rs-fMRI group average cohort consists of 1003 individuals, 998 of whom are also in the dMRI dataset. These datasets include some twin and non-twin siblings. However, individuals’ family structure, as well as exact age, handedness, and ethnicity are access-restricted to protect the privacy of the subjects and these data were not requested as they are not critical to this study. Group-average dense T1w/T2w myelination index were gathered from the same source. Macaque retrograde tracer connectivity was sourced from supplementary table 6 of (Markov et al., 2014). Parcel-by-parcel values were averaged across monkey and hemisphere. Group average, parcellated cortco-cortico evoked potential (CCEP) connectivity was gathered from the v1903 release of the Functional Brain Tractography project (F-TRACT) (David et al., 2013; Trebaul et al., 2018) at https://f-tract.eu.

### Cortical parcellation & functional networks

The HCP multimodal parcellation scheme (HCP-MMP1.0), consisting of 180 cortical parcels per hemisphere, was projected from the Workbench (Marcus et al., 2011) 32k grayordinate template brain to the FreeSurfer (Fischl, 2012) ico5 fsaverage template as per (Coalson et al., 2016). Using the FreeSurfer reconstruction directories gathered from the database, surface-based fsaverage parcel labels were mapped onto each individual’s white matter surface using spherical landmark registration (`fs_label2label`), (Fischl et al., 1999). Grayordinate rs-fMRI connectivity values were morphed to the ico5 fsaverage template then averaged within each parcel. Finally, individual’s surface-based parcel labels were converted to binary volumes marking the gray matter — white matter boundary (`mri_label2vol`) to serve as seed and target regions for probabilistic tractography. Workbench and FreeSurfer functions were sourced from releases 1.2.3 and 6.0, respectively.

To facilitate interpretation of the connectome, parcels were ordered and grouped into functional networks adapted from (Ji et al., 2019), which applied iterative Louvain clustering (Blondel et al., 2008; Rubinov and Sporns, 2010) and other criteria to a resting-state fMRI connectivity. These functional groupings and parcel order were selected as they were also generated using (a subset of) the WU-Minn HCP dataset and the HCP-MMP1.0 parcellation scheme. For this study the parcels of the left and right hemispheres were separated and the order and groupings of the left hemisphere in (Ji et al., 2019) were used for homologous parcels in the both right and left hemisphere, respectively, when combining data across hemispheres. Two pairs of the original networks (primary and secondary visual, ventral and posterior multimodal) contained too few parcels for effective analysis and were highly inter-related. These network pairs were simplified by combining them into visual and multimodal groups, yielding 10 functional networks per hemisphere, see **table 2**.

**Table 1.** Connectome Features

| Connectome Features |
| --- |
| Probabilistic methodology sensitive to weak connections yielding a fully-populated, un-thresholded connectome |
| Cortex parcellated into the standardized, relatively dense, and functionally relevant HCP-MMP1.0 atlas |
| Large normative sample size (N = 1,065) |
| Enables comparison with other measures in the WU-Minn HCP and other cohorts |

**Table 2.** Parcel order and network assignment. The emboldened indices refer to the parcel order in figure 1A. The Orig. indices refer to the original parcel order presented in (Glasser et al., 2016). All indices refer to the left hemisphere, adding 180 yields the homologous right hemisphere indices.

| **Idx.** | **Parcel** | **Orig.** | **Network** | **Idx.** | **Parcel** | **Orig.** | **Network** | **Idx.** | **Parcel** | **Orig.** | **Network** |
| --- | --- | --- | --- | --- | --- | --- | --- | --- | --- | --- | --- |
| 1 | V1 | 1 | Cingulo-Opercular | 61 | 46 | 84 | Cingulo-Opercular | 121 | IP1 | 145 | Frontoparietal |
| 2 | ProS | 121 | Visual | 62 | 9-46d | 86 | Cingulo-Opercular | 122 | PFm | 149 | Frontoparietal |
| 3 | DVT | 142 | Visual | 63 | 43 | 99 | Cingulo-Opercular | 123 | p10p | 170 | Frontoparietal |
| 4 | MST | 2 | Visual | 64 | PFcm | 105 | Cingulo-Opercular | 124 | p47r | 171 | Frontoparietal |
| 5 | V6 | 3 | Visual | 65 | Pol2 | 106 | Cingulo-Opercular | 125 | A1 | 24 | Auditory |
| 6 | V2 | 4 | Visual | 66 | FOP4 | 108 | Cingulo-Opercular | 126 | 52 | 103 | Auditory |
| 7 | V3 | 5 | Visual | 67 | MI | 109 | Cingulo-Opercular | 127 | RI | 104 | Auditory |
| 8 | V4 | 6 | Visual | 68 | FOP1 | 113 | Cingulo-Opercular | 128 | TA2 | 107 | Auditory |
| 9 | V8 | 7 | Visual | 69 | FOP3 | 114 | Cingulo-Opercular | 129 | PBelt | 124 | Auditory |
| 10 | V3A | 13 | Visual | 70 | PFop | 147 | Cingulo-Opercular | 130 | MBelt | 173 | Auditory |
| 11 | V7 | 16 | Visual | 71 | PF | 148 | Cingulo-Opercular | 131 | LBelt | 174 | Auditory |
| 12 | IPS1 | 17 | Visual | 72 | Pol1 | 167 | Cingulo-Opercular | 132 | A4 | 175 | Auditory |
| 13 | FFC | 18 | Visual | 73 | FOP5 | 169 | Cingulo-Opercular | 133 | 7m | 30 | Default Mode |
| 14 | V3B | 19 | Visual | 74 | PI | 178 | Cingulo-Opercular | 134 | POS1 | 31 | Default Mode |
| 15 | LO1 | 20 | Visual | 75 | a32pr | 179 | Cingulo-Opercular | 135 | 23d | 32 | Default Mode |
| 16 | LO2 | 21 | Visual | 76 | p24 | 180 | Cingulo-Opercular | 136 | v23ab | 33 | Default Mode |
| 17 | PIT | 22 | Visual | 77 | PEF | 11 | Dorsal Attention | 137 | d23ab | 34 | Default Mode |
| 18 | MT | 23 | Visual | 78 | 7PL | 46 | Dorsal Attention | 138 | 31pv | 35 | Default Mode |
| 19 | LIPv | 48 | Visual | 79 | MIP | 50 | Dorsal Attention | 139 | a24 | 61 | Default Mode |
| 20 | VIP | 49 | Visual | 80 | LIPd | 95 | Dorsal Attention | 140 | d32 | 62 | Default Mode |
| 21 | PH | 138 | Visual | 81 | 6a | 96 | Dorsal Attention | 141 | p32 | 64 | Default Mode |
| 22 | V6A | 152 | Visual | 82 | PFt | 116 | Dorsal Attention | 142 | 10r | 65 | Default Mode |
| 23 | VMV1 | 153 | Visual | 83 | AIP | 117 | Dorsal Attention | 143 | 47m | 66 | Default Mode |
| 24 | VMV3 | 154 | Visual | 84 | PHA3 | 127 | Dorsal Attention | 144 | 8Av | 67 | Default Mode |
| 25 | V4t | 156 | Visual | 85 | TE2p | 136 | Dorsal Attention | 145 | 8Ad | 68 | Default Mode |
| 26 | FST | 157 | Visual | 86 | PHT | 137 | Dorsal Attention | 146 | 9m | 69 | Default Mode |
| 27 | V3CD | 158 | Visual | 87 | PGp | 143 | Dorsal Attention | 147 | 8BL | 70 | Default Mode |
| 28 | LO3 | 159 | Visual | 88 | IP0 | 146 | Dorsal Attention | 148 | 9p | 71 | Default Mode |
| 29 | VMV2 | 160 | Visual | 89 | 55b | 12 | Language | 149 | 10d | 72 | Default Mode |
| 30 | VVC | 163 | Visual | 90 | PSL | 25 | Language | 150 | 47l | 76 | Default Mode |
| 31 | 4 | 8 | Somatomotor | 91 | SFL | 26 | Language | 151 | 9a | 87 | Default Mode |
| 32 | 3b | 9 | Somatomotor | 92 | STV | 28 | Language | 152 | 10v | 88 | Default Mode |
| 33 | 5m | 36 | Somatomotor | 93 | 44 | 74 | Language | 153 | 10pp | 90 | Default Mode |
| 34 | 5L | 39 | Somatomotor | 94 | 45 | 75 | Language | 154 | OFC | 93 | Default Mode |
| 35 | 24dd | 40 | Somatomotor | 95 | IFJa | 79 | Language | 155 | 47s | 94 | Default Mode |
| 36 | 24dv | 41 | Somatomotor | 96 | IFSp | 81 | Language | 156 | EC | 118 | Default Mode |
| 37 | 7AL | 42 | Somatomotor | 97 | STGa | 123 | Language | 157 | PreS | 119 | Default Mode |
| 38 | 7PC | 47 | Somatomotor | 98 | A5 | 125 | Language | 158 | H | 120 | Default Mode |
| 39 | 1 | 51 | Somatomotor | 99 | STSda | 128 | Language | 159 | PHA1 | 126 | Default Mode |
| 40 | 2 | 52 | Somatomotor | 100 | STSdp | 129 | Language | 160 | STSvp | 130 | Default Mode |
| 41 | 3a | 53 | Somatomotor | 101 | TPOJ1 | 139 | Language | 161 | TGd | 131 | Default Mode |
| 42 | 6d | 54 | Somatomotor | 102 | TGv | 172 | Language | 162 | TE1a | 132 | Default Mode |
| 43 | 6mp | 55 | Somatomotor | 103 | RSC | 14 | Frontoparietal | 163 | TE2a | 134 | Default Mode |
| 44 | 6v | 56 | Somatomotor | 104 | POS2 | 15 | Frontoparietal | 164 | PGi | 150 | Default Mode |
| 45 | OP4 | 100 | Somatomotor | 105 | 7Pm | 29 | Frontoparietal | 165 | PGs | 151 | Default Mode |
| 46 | OP1 | 101 | Somatomotor | 106 | 8BM | 63 | Frontoparietal | 166 | PHA2 | 155 | Default Mode |
| 47 | OP2-3 | 102 | Somatomotor | 107 | 8C | 73 | Frontoparietal | 167 | 31pd | 161 | Default Mode |
| 48 | FOP2 | 115 | Somatomotor | 108 | a47r | 77 | Frontoparietal | 168 | 31a | 162 | Default Mode |
| 49 | Ig | 168 | Somatomotor | 109 | IFJp | 80 | Frontoparietal | 169 | 25 | 164 | Default Mode |
| 50 | FEF | 10 | Cingulo-Opercular | 110 | IFSa | 82 | Frontoparietal | 170 | s32 | 165 | Default Mode |
| 51 | 5mv | 37 | Cingulo-Opercular | 111 | p9-46v | 83 | Frontoparietal | 171 | STSva | 176 | Default Mode |
| 52 | 23c | 38 | Cingulo-Opercular | 112 | a9-46v | 85 | Frontoparietal | 172 | TE1m | 177 | Default Mode |
| 53 | SCEF | 43 | Cingulo-Opercular | 113 | a10p | 89 | Frontoparietal | 173 | PCV | 27 | Multimodal |
| 54 | 6ma | 44 | Cingulo-Opercular | 114 | 11l | 91 | Frontoparietal | 174 | TPOJ2 | 140 | Multimodal |
| 55 | 7Am | 45 | Cingulo-Opercular | 115 | 13l | 92 | Frontoparietal | 175 | TPOJ3 | 141 | Multimodal |
| 56 | p24pr | 57 | Cingulo-Opercular | 116 | i6-8 | 97 | Frontoparietal | 176 | PeEc | 122 | Multimodal |
| 57 | 33pr | 58 | Cingulo-Opercular | 117 | s6-8 | 98 | Frontoparietal | 177 | TF | 135 | Multimodal |
| 58 | a24pr | 59 | Cingulo-Opercular | 118 | AVI | 111 | Frontoparietal | 178 | Pir | 110 | Orbito-Affective |
| 59 | p32pr | 60 | Cingulo-Opercular | 119 | TE1p | 133 | Frontoparietal | 179 | AAIC | 112 | Orbito-Affective |
| 60 | 6r | 78 | Cingulo-Opercular | 120 | IP2 | 144 | Frontoparietal | 180 | pOFC | 166 | Orbito-Affective |

**Table 3.** Statistics and uncertainty. Where multiple uncertainties are listed for a figure panel, they correspond to the statistics read left-to-right, top-to-bottom in that panel. For figure 8B only uncertainties for significant correlations are listed. Uncertainties for figures 6, 7, 8 and 10 are not shown. Figure 6-1 contains bootstrapped 95% confidence intervals for the 180 means shown in figure 6, n = 179. Figure 7 shows Bootstrapped 95% confidence intervals in gray; the values of these intervals for all distance bins are available in the figure source data at https://doi.org/10.5281/zenodo.4060485. For figure 10 means across shuffled matrices are only necessary to account for arbitrary ordering among tied edge weights and the bootstrapped 95% confidence intervals for these means are vanishingly small. The values of these intervals at all network densities are also included in the figure source data. For nonlinear regressions confidence intervals are estimated using *R*^-1^, the inverse *R* factor from *QR* decomposition of the Jacobian, the degrees of freedom for error, and the root mean squared error. For linear correlations the confidence intervals are based on an asymptotic normal distribution of 0.5*log((1+r)/(1-r)), with an approximate variance equal to 1/(N-3). For descriptive statistics, e.g. means, empirical 95% confidence intervals are estimated by bootstrapping with 2000 iterations.

| Location | Data structure | Test or analysis | N | Uncertainty [CI<sub>95%</sub>] |
| --- | --- | --- | --- | --- |
| Fig 1-1D | Gaussian predictor<br>Exponential response | Nonlinear regression<br>(iterative optimization) | 64,620 | $\lambda = 23.8$ [23.5 24.0] |
| | | | 64,620 | $\lambda = 22.8$ [22.7 22.9] |
| | | | 64,620 | $\lambda = 22.2$ [22.1 22.2] |
| | | | 64,620 | $\lambda = 23.4$ [23.3 23.6] |
| Fig 2A | Gaussian predictor<br>Exponential response | Nonlinear regression<br>(iterative optimization) | 16,110 | $\lambda = 23.1$ [22.8 23.3] |
| Fig 2B | Gaussian predictor<br>Exponential response | Nonlinear regression<br>(iterative optimization) | 16,110 | $\lambda = 23.9$ [23.7 24.2] |
| Fig 2C | Gaussian predictor<br>Exponential response | Nonlinear regression<br>(iterative optimization) | 32,400 | $\lambda = 32.8$ [32.5 33.0] |
| Fig 2D | Gaussian predictor<br>Exponential response | Nonlinear regression<br>(iterative optimization) | 64,620 | $\lambda = 23.4$ [23.3 23.6] |
| Fig 2-2B | Gaussian predictor<br>Gaussian response | Linear correlation | 1,065 | $r = -0.14$ [-0.20 -0.08] |
| Fig 3D | Gaussian predictor<br>Exponential response | Nonlinear regression<br>(iterative optimization) | 12,924 | $\lambda = 27.8$ [27.4 28.2] |
| Fig 3F | Gaussian predictor<br>Gaussian response | Linear correlation | 1,065 | $r = 0.70$ [0.67 0.73] |
| Fig 4C | Gaussian predictor<br>Gaussian response | Linear correlation | 80 | $r = 0.35$ [0.14 0.53] |
| Fig 8A | Gaussian predictor<br>Gaussian response | Linear correlation | 16,110 | $r = -0.10$ [-0.12 -0.09] |
| | | | 16,110 | $r = -0.12$ [-0.13 -0.10] |
| | | | 32,400 | $r = -0.11$ [-0.12 -0.10] |
| Fig 8B | Gaussian predictor<br>Gaussian response | Linear correlation | 351 | $r = -0.17$ [-0.27 -0.06] |
| | | | 351 | $r = -0.13$ [-0.23 -0.02] |
| | | | 66 | $r = -0.41$ [-0.60 -0.19] |
| | | | 91 | $r = -0.26$ [-0.44 -0.06] |
| | | | 231 | $r = -0.30$ [-0.42 -0.18] |
| | | | 231 | $r = -0.30$ [-0.40 -0.17] |
| | | | 28 | $r = -0.56$ [-0.77 -0.24] |
| | | | 780 | $r = -0.12$ [-0.19 -0.05] |
| | | | 780 | $r = -0.17$ [-0.24 -0.10] |
| | | | 10 | $r = -0.74$ [-0.93 -0.20] |
| Fig 9B | Gaussian predictor<br>Gaussian response | Linear correlation | 19,667 | $r = 0.43$ [0.42 0.44] |
| | | | 19,667 | $r = 0.23$ [0.21 0.24] |
| | | | 64,620 | $r = 0.06$ [0.05 0.07] |
| Figure 9-1 | Gaussian predictor<br>Gaussian response | Linear correlation | 8,483 | $r = 0.42$ [0.40 0.44] |
| | | | 8,483 | $r = 0.22$ [0.20 0.24] |
| | | | 16,110 | $r = 0.06$ [0.05 0.07] |
| | | | 8,370 | $r = 0.40$ [0.38 0.42] |
| | | | 8,370 | $r = 0.22$ [0.20 0.24] |
| | | | 16,110 | $r = 0.11$ [0.10 0.13] |

### Probabilistic tractography

All analysis of diffusion imaging data was performed with FSL (Behrens et al., 2007; Jenkinson et al., 2012) release 6.0.1. Analyses were performed identically for each subject and broadly follow (Burns, 2014). The diffusion and bedpostX precursor directories made available from the HCP database were used as inputs without modification. The WU-Minn HCP diffusion data are corrected for eddy currents and movement with FSL eddy (Andersson and Sotiropoulos, 2016). Subjects’ estimated displacement over time from their initial position is written to the eddy_restricted_movement_rms output. Using these data, a scalar index of each subject’s motion was derived by integrating their displacement over time.

Fractional anisotropy (FA) analysis was performed using `dtifit`. The resulting FA volumes were not analyzed but only used for registering the FreeSurfer and dMRI volumes (`flirt`), as is necessary to map the parcel masks into dMRI space (`probtrackx2` arguments `--xfm -- seedref`). Non-invasive probabilistic tractography was performed with `probtrackx2` in voxel-by-parcel mode (`--os2t --s2tastext`). In this configuration, the number and length of streamlines (`--ompl --opd`) is estimated from each voxel in the seed parcel to each target parcel as a whole. To aid parallelization of these computationally intensive processes, the list of target parcels (`--targetmasks`) was quartered into four sub-lists. Therefore `probtrackx2` was invoked 1440 times per subject, estimating the connectivity between 1 seed parcel and 90 target parcels in each invocation. The default ½ voxel step length, 5000 samples and 2000 steps were used (`--steplength 0.5 -P 5000 -S 2000`). To avoid artifactual loops, streamlines that loop back on themselves were discarded (`-l`) and tractography was constrained by a 90° threshold (`-c 0`) for maximal curvature between successive steps. Within-parcel connectivity and cotico-subcortical connectivity were not examined in this study. All post-hoc analyses and visualization of connectivity data were performed in Matlab 2019b (Mathworks) except for **figure 1C** which was rendered in fsleyes.

**Figure 1.**
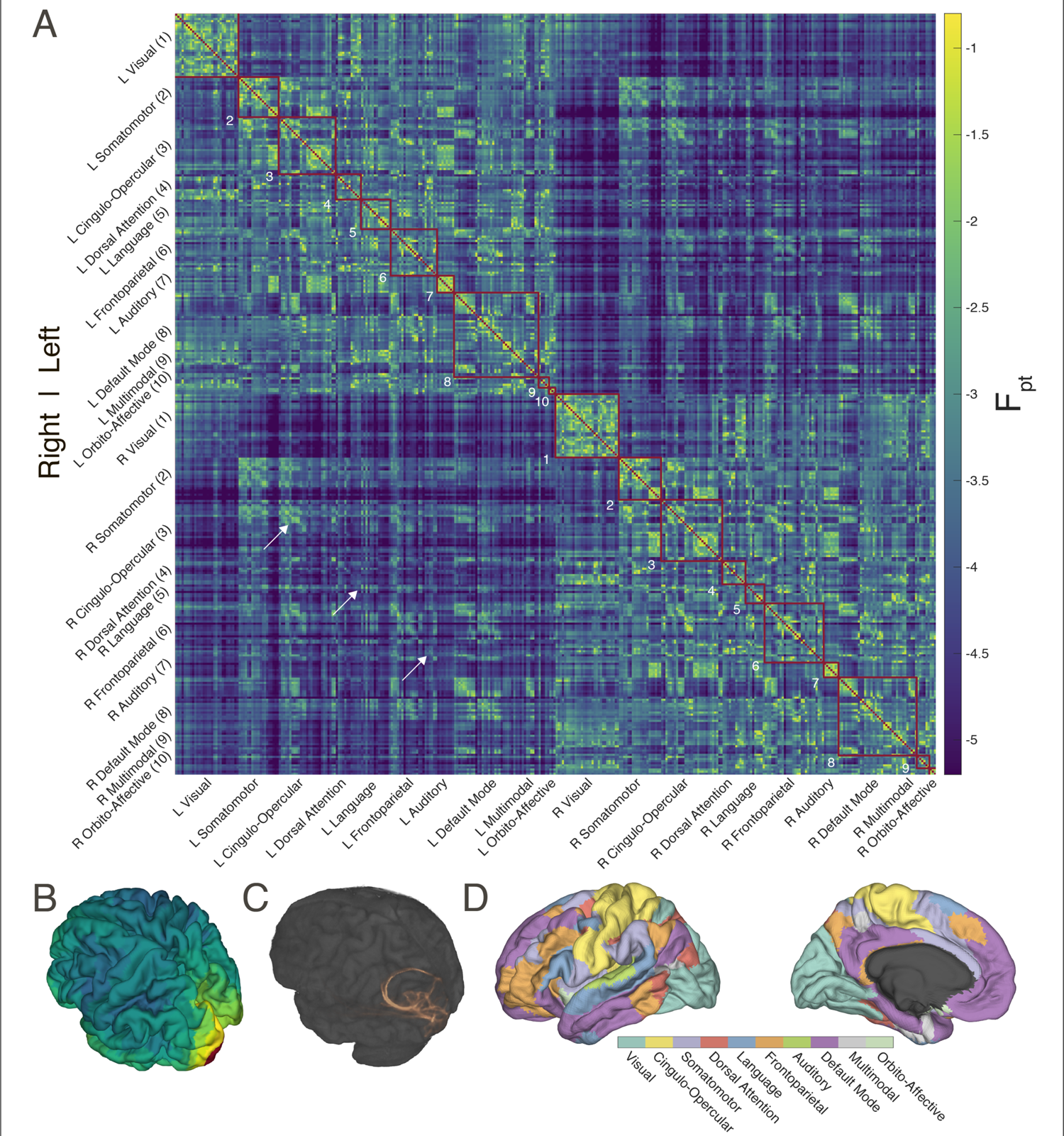
Probabilistic diffusion tractography structural connectome of the human cortex (**A**) Group average (N = 1065) structural connectivity matrix consisting of the 360 HCP-MMPS1.0 atlas parcels organized into ten functional networks. Raw streamline counts are fractionally scaled yielding the log probability Fpt. The white arrows highlight the diagonal which contains contralateral homologs. (**B**) The first row of the connectivity matrix, showing connection probabilities from left V1 to all other parcels, projected onto the fsaverage template cortex. (**C**) Single subject (100307) volume ray casting visualization of left V1-originating streamline probabilities within the skull-stripped T1-weighted structural MR volume. (**D**) Ten functional networks, adapted from (Ji et al., 2019), within HCP-MMPS1.0 atlas. These are indicated by red boxes in panel **A**.

**Figure 1-1.**
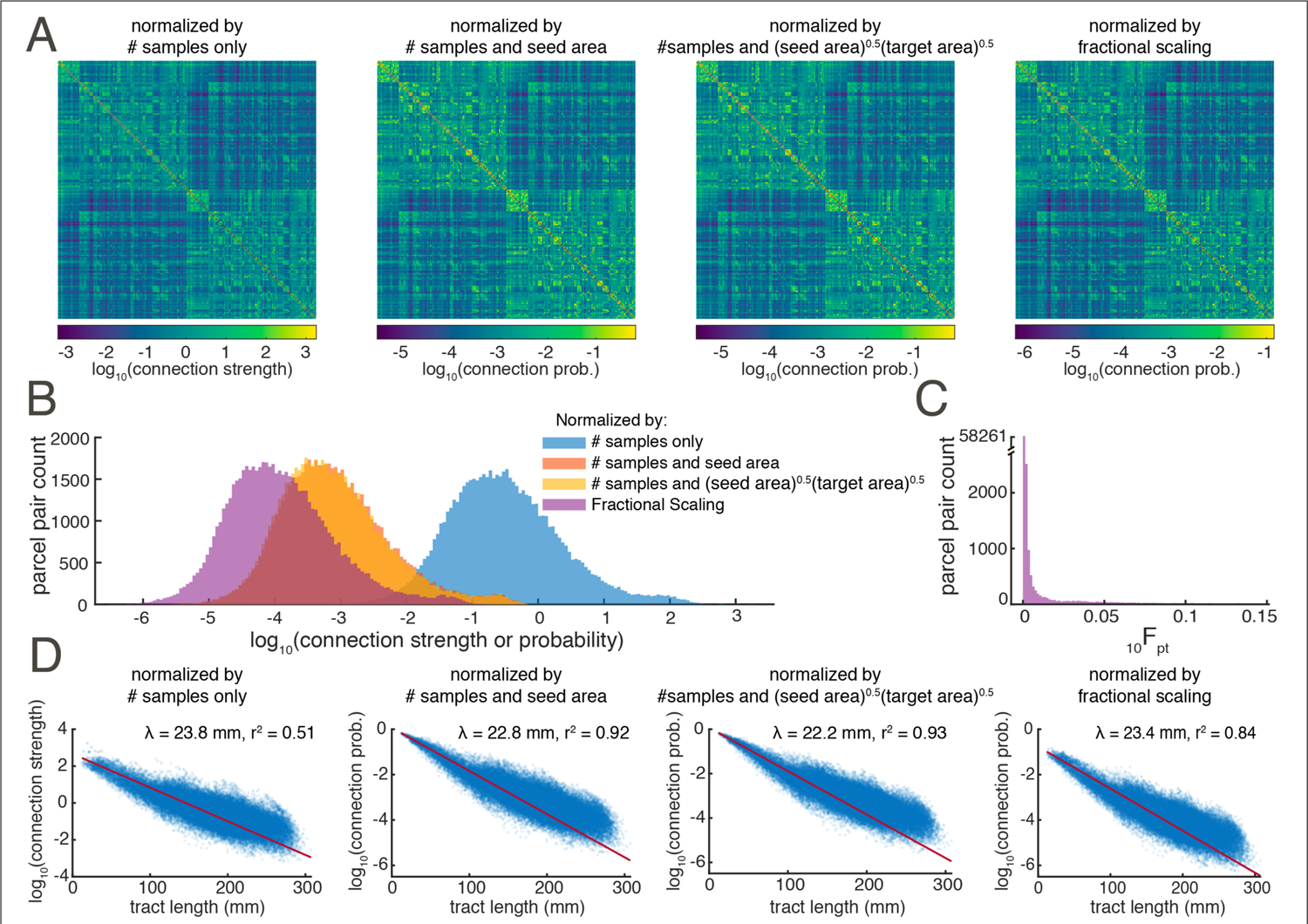
Comparison of normalization methods. Shown are the (**A**) connectivity matrices, (**B**) distributions of pairwise connectivity, (**C**) the pre-log distribution of Fpt (**D**) relationships between connectivity and fiber tract length for four normalization methods.

### Normalization & symmetrization

Raw streamline counts were averaged across all subjects, then normalized and symmetrized following procedure developed for non-human primate histological tracing (Donahue et al., 2016; Theodoni et al., 2020). Briefly, fractionally scaled values are defined as the ratio of the number of streamlines originating at parcel A and terminating at parcel B to the total number of streamlines that either originate at parcel A or terminate at parcel B while excluding within-parcel connections.

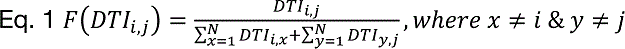

Fractional scaling is one of several plausible normalization strategies. Because we used 5000 samples (-P 5000) and voxel-by-parcel mode (--os2t) in our probtrackX invocation, the maximum possible raw streamline count between any two parcels is 5000N where N is the # of voxels in the seed parcel. Note that because, for probtrackX, all parcels were defined as a single layer of 1mm isotropic voxels at the white matter — gray matter interface, Ni is also equivalent to the area of the seed parcel, in mm^2^. As shown in extended data **figure 1-1**, We examined four strategies for normalizing the raw streamline counts: (1) dividing by the number of samples, 5000, (2) dividing by the number of samples and seed area, 5000Ni, (3) dividing by the number of samples and the areas of both the seed and target parcels, 5000N_i_^0.5^N_j_^0.5^, and (4) fractional scaling, see Eq. 1. These approaches yield similar connectivity matrices, distributions of pairwise connectivity, and rates of connectivity fall-off with fiber tract distance. The choice of normalization does shift the absolute scale of pairwise connectivity strengths, but as this effect is mostly homogenous across all connections, subsequent analyses are not greatly affected.

The correlation coefficient of connectivity strengths between normalization techniques exceeds 0.97 for all pairwise comparisons, and exceeds 0.99 if the samples-only normalization approach is excluded (data not shown).

While diffusion tractography is not sensitive to the directionality of connections, because parcel A to B and parcel B to A streamlines are computed separately minor asymmetries arise. Connectivity matrix symmetry is enforced by taking the arithmetic mean of the A-B and B-A fractionally scaled connection weights.

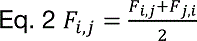

Because probabilistic tractography values span several orders of magnitude, and are approximately log-normally distributed (**Fig. 1-1 B**), data were log-transformed (log_10_) prior to subsequent analyses. The CCEP and rs-fMRI connectivity matrices were (re)normalized following the same procedure. However the rsMRI connectivity values were not log-transformed because these data are already approximately normally distributed, if bimodal, in linear space, see **figure 9B**.

### Network theory metrics

All network theoretic measures were computed in matlab using the Brain Connectivity Toolbox, 2019-03-03 release (Rubinov and Sporns, 2010). It is available at http://www.brain-connectivity-toolbox.net or https://www.nitrc.org/projects/bct. The definitions for the metrics used (for binary and undirected networks) are repeated below.

### Precursor measures

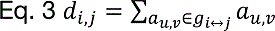

Where *d_i,j_* is the shortest path length, a basis for measuring integration, between nodes *i* and *j*, *N* is the set of all nodes in the network, *n* is the number of nodes, and *au,v* is the binarized connectivity between nodes *u* and *v*.

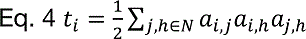

Where *t_i_* is the number of triangles, a basis for measuring integration, around node *i*.

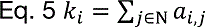

Where *k_i_* is the number of degree, or number of links, connected to node *i*.

### Mean Clustering Coefficient (MCC)

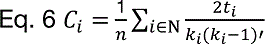

Where *C_i_* is the clustering coefficient of node *i*. (*C_i_* = 0 for *k_i_* < 2), (Watts and Strogatz, 1998).

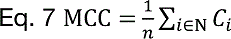

### Characteristic Path Length (CPL)

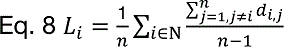

Where *L_i_* is the number of the average distance between node *i* and all other nodes, (Watts and Strogatz, 1998).

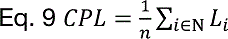

### Global Efficiency

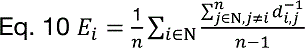

Where *Ei* is the efficiency of node *i*.

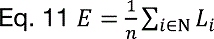

Where *E* is the global efficiency of the network, (Latora and Marchiori, 2001).

### Modularity

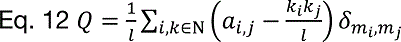

Where *l* is the number of links in the network, *m_i_* is module containing node *i*, *δ_mi,mj_* = 1 if *m_i_* = *mj*, and 0 otherwise, and *Q* is the global efficiency of the network, (Newman, 2004).

### Gamma (normalized MCC)

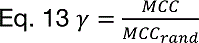

Where *MCC_rand_* is the *MCC* of a random network of the same statistical makeup.

### Lambda (normalized CPL)

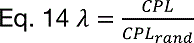

Where *CPL_rand_* is the *CPL* of a random network of the same statistical makeup. Note that this measure is unrelated to the length constant λ.

### Small Worldness

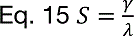

Where *S* is the network small-worldness (Humphries and Gurney, 2008).

### Transitivity

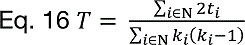

Where *T* is the transitivity of the network (Newman, 2003).

### Assortativity

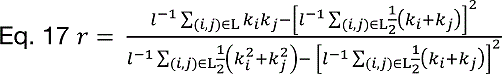

Where *L* is the set of all links and *r* is the assortativity coefficient of the network (Newman, 2003).

### Network Density

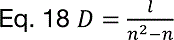

Where *D* is the density of the network before thresholding and binarization.

## Results

### A whole-cortex structural connectome

**Figure 1A** shows the group average parcel to parcel and probabilistic diffusion tractography connectome. This matrix consists of connectivity among the 360 cortical parcels of the HCP-MMP1.0 atlas. Using left V1 connectivity as an example, **figures 1B** illustrates the spatial mapping of the connectivity matrix to the cortex, and **1C** shows a rendering of streamline paths for one subject. The cortical parcels are further organized into 10 functional groups per hemisphere modified from (Ji et al., 2019). These larger functional groupings are shown in **figure 1D**. The raw probabilistic tractography streamline counts have been normalized by fractionally scaling (Eq. 1) into log probabilities (F_pt_) following procedures developed for tracing non-human primate connectivity. As dMRI reveals structural connections, the network is undirected and therefore symmetric. The main diagonal is masked as intra-parcel connectivity was not examined in this study. The upper left quadrant shows connectivity among the 180 parcels of the left hemisphere, the lower right quadrant the connectivity within the right hemisphere. The upper right and lower left quadrants are duplicates and show the inter-hemispheric, or callosal, connections. The 180^th^ (or half-) diagonal is clearly visible (white arrows); this shows the connectivity between homologous parcels in the right and left hemispheres, which is greater than non-homologous callosal connectivity for most parcels.

After log_10_ transformation, F_pt_ connectivity among all parcel pairs is approximately Gaussian in distribution with mean −3.903 (CI_95%_ = [−3.910 −3.897]), standard deviation 0.8111 (CI_95%_ = [0.806 0.816]), skewness 0.627 (CI_95%_ = [0.602 0.644]), and kurtosis 3.605 (CI_95%_ = [3.560 3.650]). In addition to bringing the range of F_pt_ values into the same order of magnitude, log_10_ transformation is justified as it brings the distribution’s skewness significantly closer to zero (pre-log_10_: 9.047, CI_95%_ = [8.719 9.469]), and kurtosis significantly closer to three, pre-log_10_: 103.684 (CI_95%_ = [93.991 117.026]) thus bringing the distribution closer to normality. See extended data **figure 1-1 B, C** for a graphical comparison. Empirical confidence intervals were estimated via bootstrapping with 2000 iterations. The values of the group average and individual probabilistic dMRI connectivity matrices, as well as all other figure source data can be found at https://doi.org/10.5281/zenodo.4060485.

### Tract length strongly predicts connectivity strength, with exponential decay

In addition to the connection strength, diffusion tractography estimates the fiber tract length between all pairs of parcels. As shown in **figure 2**, structural connectivity (10^F_pt_) falls off as an exponential function of fiber tract length with the form 10^F_pt_ = α*e^-d/λ^ where λ is the length constant, α the scaling coeffect, and d the tract length. Alternative functional forms were examined (see **figure 2-1**), but the exponential was selected for parsimony, goodness-of-fit, and concordance with histological tracing data (see Discussion). Note that λ is sometimes reported in inverted units of mm^-1^, e.g. (Markov et al., 2013; Theodoni et al., 2020), but we here use the λ convention from neuronal cable theory (Dayan and Abbott, 2001) which has more intuitive units (mm); the conventions are conceptually equivalent. For the group-average connectome, λ = 23.4 mm and the least squares exponential fit explains 84% of the variance in 10^F_pt_ across all parcel pairs. Callosal connectivity, when isolated, decays more slowly with respect to tract length, λ = 32.8, and hews to the exponential expectation less consistently r^2^ = 0.62. Because the tracing of long fiber tracts may be hampered by poor scan quality, we investigated the effects of subjects’ motion on λ. For each subject, λ was calculated for non-zero connections in the same manner as the group average. While subjects’ motion within the scanner does reduce λ, this effect is modest, only explaining 1.96% of the inter-subject variance, see **figure 2-2**.

**Figure 2.**
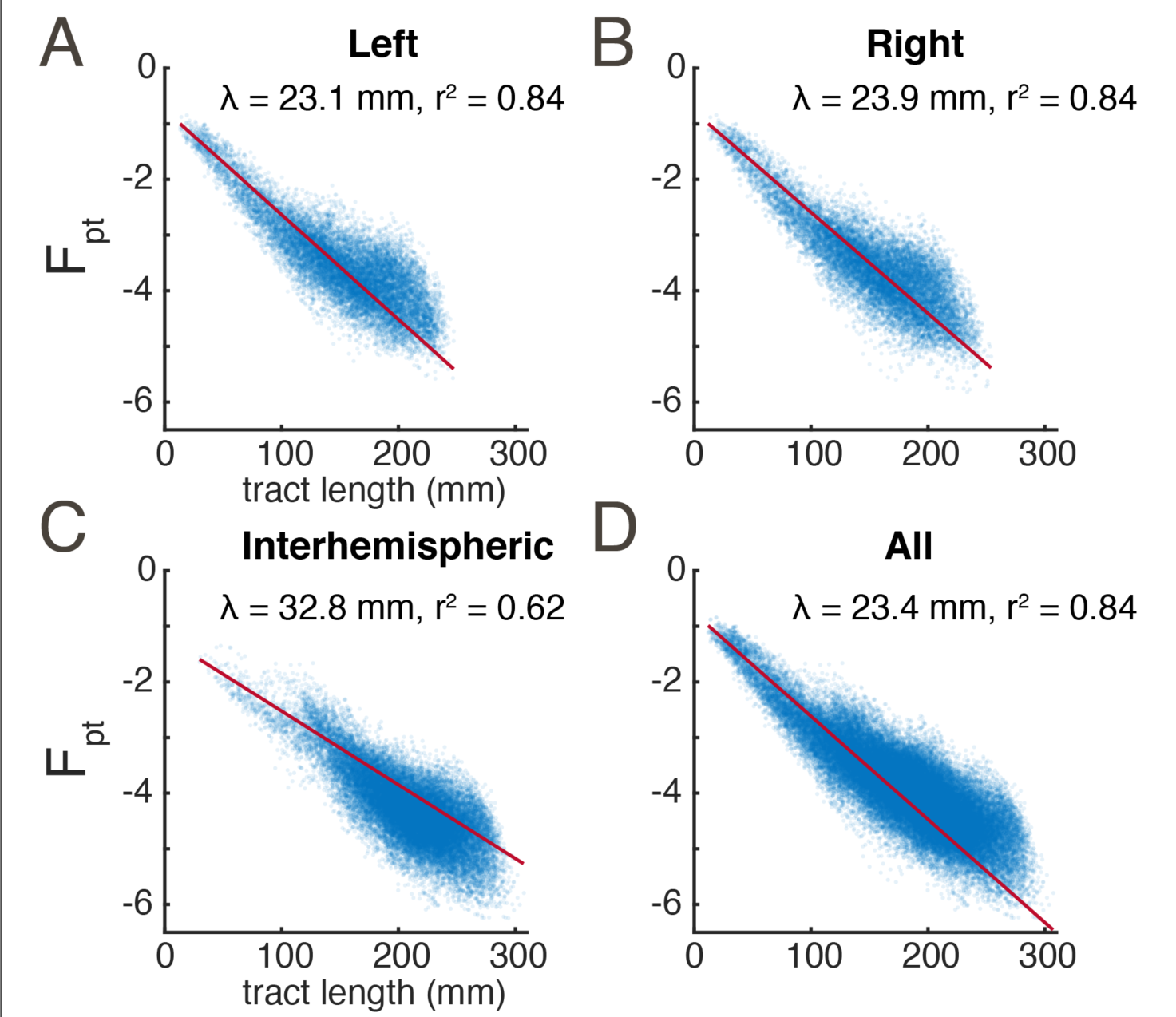
Connectivity strength exponential decays with fiber tract length. (**A**) and (**B**) connections within the right and left hemispheres, respectively. (**C**) Connections between the right and left hemisphere. (**D**) All connections. Each marker represents a pair of parcels. Red traces show the least-squares exponential fit; inset are the length constant λ and r^2^ of this fit. Note that F_pt_ is log-transformed making these axes effectively semi-log.

**Figure 2-1.**
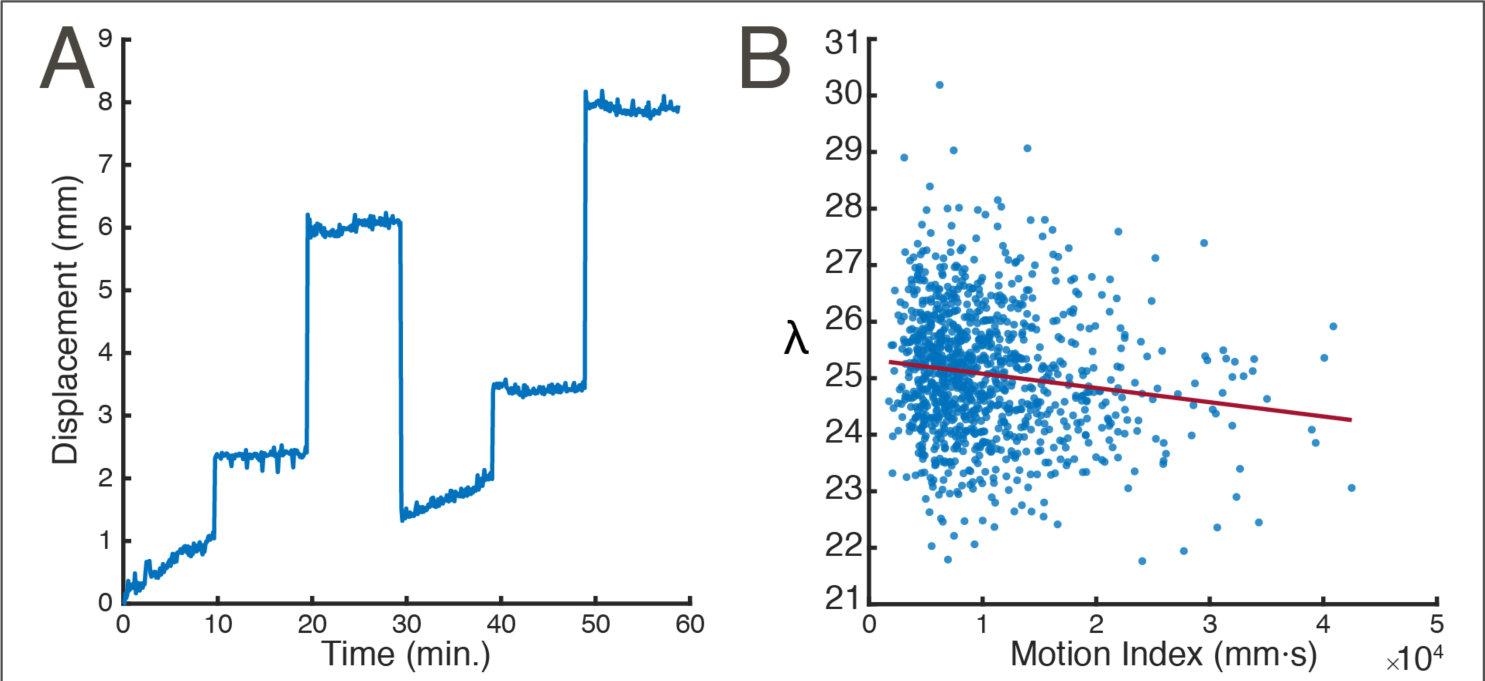
Alternative models for fitting connectivity strength as a function of fiber tract length. Each gray marker shows the average pair-wise F_pt_ between two parcels and fiber tract length between them, as also shown in figure 2D. The colored traces show maximum likelihood estimates for several listed functional forms. The AIC, AICc, aBIC columns contain the Akaike, corrected Akaike, and Bayesian information criteria, respectively. While the Gaussian fits explain slightly more variance and have a slightly lower AIC than the exponential fit, the exponential has fewer parameters and is consistent with histological non-human primate evidence (Donahue et al., 2016; Markov et al., 2013; Theodoni et al., 2020).

**Figure 2-2.**
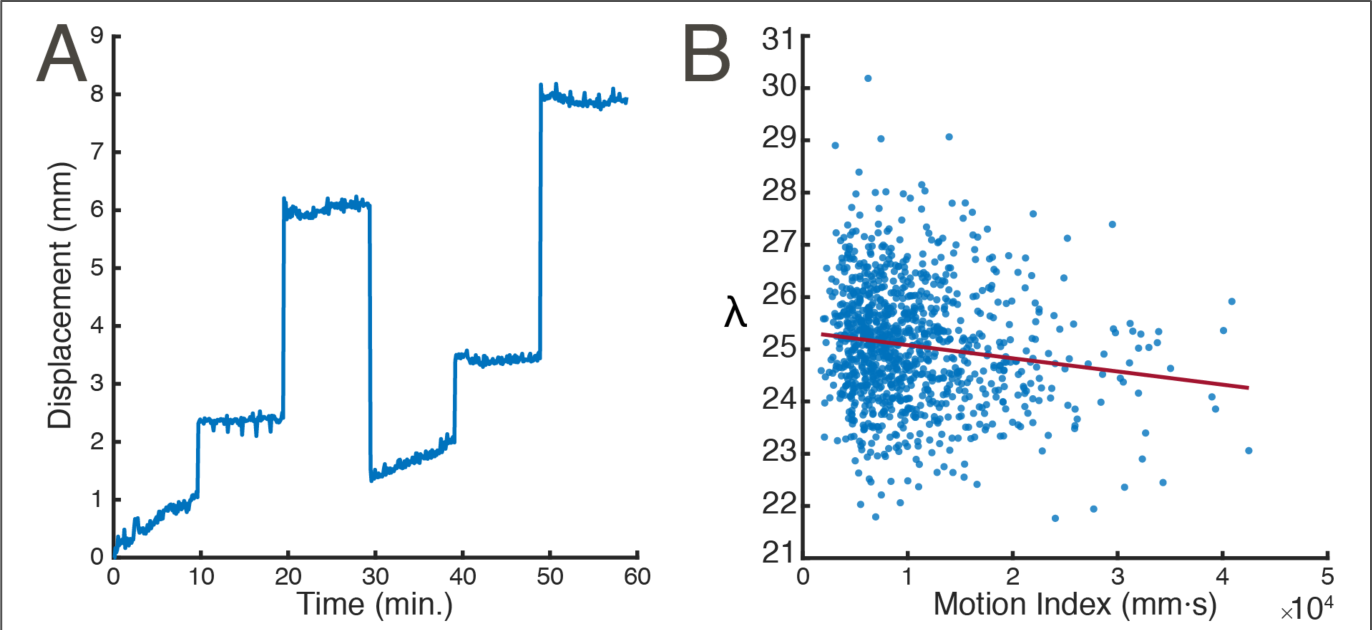
Effect of motion during the dMRI scan. (**A**) Time-course of displacement relative to initial position for one subject (996782). The six runs of the HCP dMRI protocol can be seen. (**B**) Exponential fall-off coefficient λ is only modestly affected by motion, r = 0.140, p = 4.6E-6. Each marker represents a subject.

### Inter-individual variability

The inter-individual variability of connectivity was assessed by deriving the across-subject coefficient of variation, CV, for each pairwise connection F_pt_, see **figure 3**. For this analysis, the normalization, symmetrization, and log_10_-transformation of raw connectivity values was performed on each subject. Pairwise connections with zero streamlines were not log-transformed in order to avoid infinities. While there is no clear relationship between fiber tract distance and inter-individual variability, the most consistent connection appear in two clusters of around 50-100 mm and 170-225 mm (**Fig. 3B**). When the most consistent quintile of connections is isolated (Roberts et al., 2017), connectivity falls off more slowly with tract distance, with λ increasing to ∼28 mm (**Fig. 3D**). Since the proportional size of V1/V2 varies ∼3-fold across individuals and is highly heritable (Yoon et al., 2019), we hypothesized that the ipsilateral V1-V2 connection would also be highly variable, with that variability being correlated across hemispheres. Indeed, we find that the ipsilateral V1 – V2 connection is very strong, with ∼1.8 fold variability which is strongly correlated across hemispheres (r=0.70). The scatter-plot of right vs. left F_pt_ values for this connection across subjects (**Fig. 3F**) does not reveal obvious outliers which would be indicative of subject-specific artifacts. This analysis of inter-individual variability should be considered preliminary. The WU-Minn HCP dataset is rich in individual data, including the NIH neuropsychological toolbox (Gershon et al., 2013), twin and non-twin siblings subsets, and genotypic data (<u>dbGaP phs001364.v1.p1</u>), though the latter two data types are only available by application in order to ensure subject anonymity. With access to these data, a full examination of inter-individual variability, including assessing the heritability and genetic correlates of the strength of specific connections could be made.

**Figure 3.**
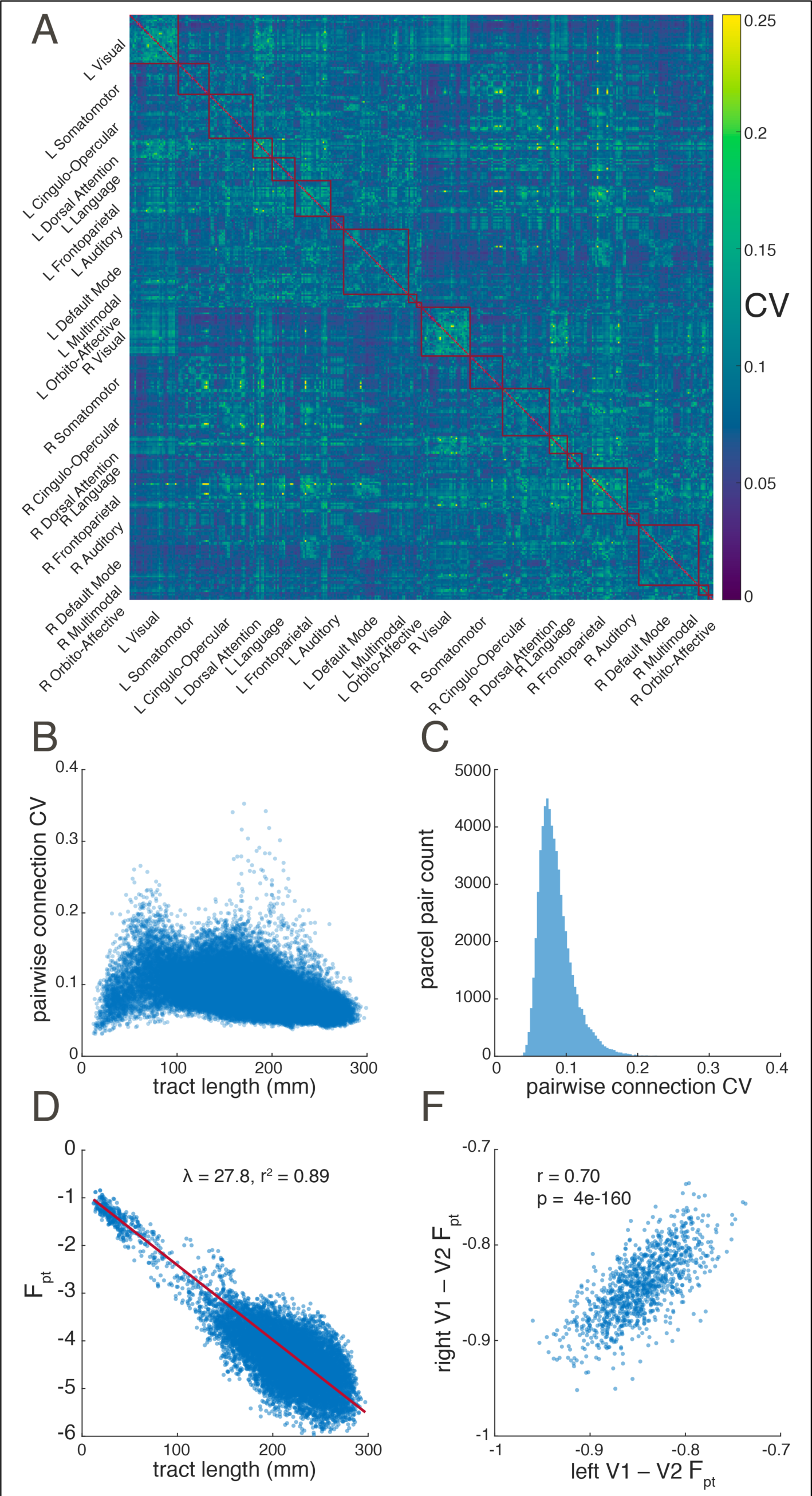
Inter-individual variability. Shown are (**A**) the matrix of connectivity coefficients of variation (CV) across subjects (**B**) pairwise CV vs. fiber tract length, (**C**) the distribution of CV across all connections, (**D**) the F_pt_ vs. fiber tract length for the connections in the highest quintile of inter-individual consistency, and (**F**) the F_pt_ of right hemisphere V1 – V2 connection in all subjects vs. left hemisphere V1 – V2 connection. In panels **B** and **D** each marker represents a sample statistic for a connection between two parcels. In panel **F** each marker represents an individual subject. In panel **D** the red trace show the least-squares exponential fit and inset are the length constant λ and r^2^ of this fit. Note that F_pt_ is log-transformed making this panel’s axes effectively semi-log. In panel F, the r^2^ of the least squares linear fit is reported.

### Probabilistic dMRI tract tracing in humans reasonably corresponds with histological fiber tracing in macaques

The development of both the HCP-MMP1.0 human cortical atlas (Glasser et al., 2016) and FV91 macaque parcellation scheme (Felleman and Van Essen, 1991) were led by David Van Essen and the parcel definitions of the human atlas were informed by human-macaque homology. As such, the parcel names of these atlases have considerable overlap, particularly for visual and visual association areas as well as the non-visual parcels 1, 2, 25, and 44. We therefore assumed that parcels with the same name were roughly homologous and limited the scope of the inter-species comparison to these parcels. Furthermore, the macaque FLne values found in (Markov et al., 2014) are directly comparable to fractionally scaled F_pt_ values (Donahue et al., 2016). Comparing the pairwise connectivity between species, we found a Pearson correlation of *r* = 0.35 (p = 0.0013), see figure 4. Considering that for macaques, Donahue and colleages (2016) found a *within-species*, between-technique correlation of *r* = 0.59 when comparing retrograde tracing and probabilistic diffusion tractography, we find the magnitude of *between-species* correlation to be reasonable supporting evidence for the efficacy of the technique.

**Figure 4.**
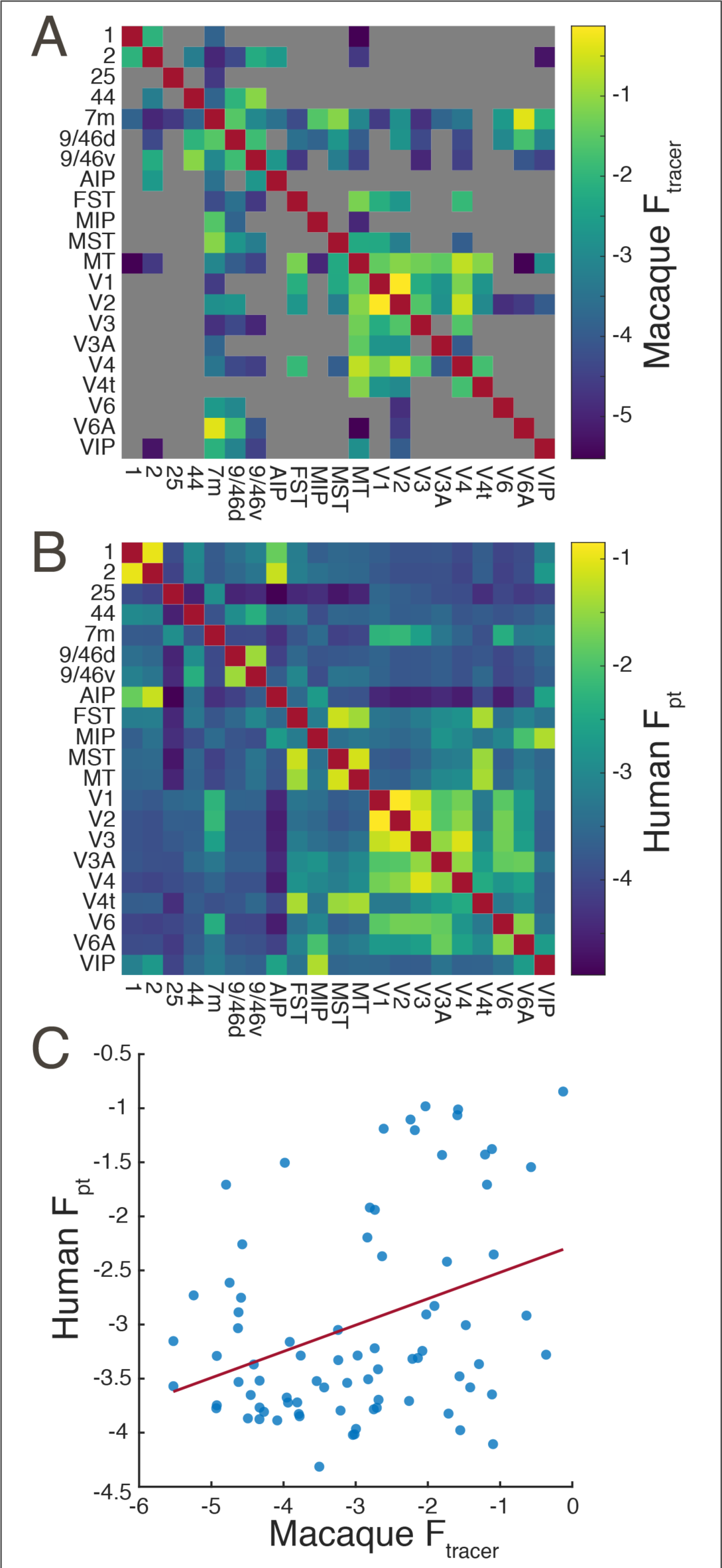
Comparison of human diffusion tractography and macaque retrograde tracing connectomes. Subset of homologous parcels in the human HCP-MMPS1.0 and macaque fv91 atlas. (**A**) Macaque group-average retrograde tracer derived structural connectome, gray indicates missing data. (**B**) Human probabilistic diffusion tractography connectome. (**C**) Pairwise correlation between macaque and human structural connectivity, r = 0.35, p = 0.0013.

### Contralateral connectivity exceeds ipsilateral connectivity in some regions

On the whole, cortical connectivity is dominated by ipsilateral connections. This effect is readily-observed by comparing the ipsilateral and contralateral quadrants of **figure 1A**.

However, there are exceptions to this rule. The differential connectome of ipsi-vs. contra-lateral connections is shown in **figure 5**. This is achieved by subtracting the mean of left-right and right-left contralateral connectivity from the mean of the right and left ipsilateral connectivity, i.e. subtracting the mean of the first and third quadrants from the mean of the second and fourth. A cingulo-parietal somatomotor region (parcels 5m, 5L, 24dd, and 24dv) are more strongly connected to most contralateral cortex than ipsilateral cortex. Lateromedial connectivity in select prefrontal (a10p, a9-46v, a10p, p10p, p47r, p9-46v, 11l, IFSa, IFJp, a24, d32, p32, 10r) and postcentral – superior parietal lobule (LIPv, VIP, 7AL, 7PC, 1, 2, 3a, 6d, 31a, 31pd, PCV) regions is stronger between hemispheres than within them. We speculate that a possible commonality between these three regions is that they have been broadly implicated in the unitary processes of somatosensory object recognition, emotion, and spatial cognition, respectively. Conversely, the entire auditory network and superior temporal cortices (STGa, STSda, DTDdp, A5, and TPOJ1) as well as the operculum and temporoparietal junction (Ig, MI, FOP1-FOP5, OP1-OP4, PF, PFcm, PFop, PI, PoI1, PoI2, and 43) have pronounced hyper-ipsilateral connectivity, consistent with the low transmission latency required for auditory processing, the left-lateralization of language, and the right lateralization of attention.

**Figure 5.**
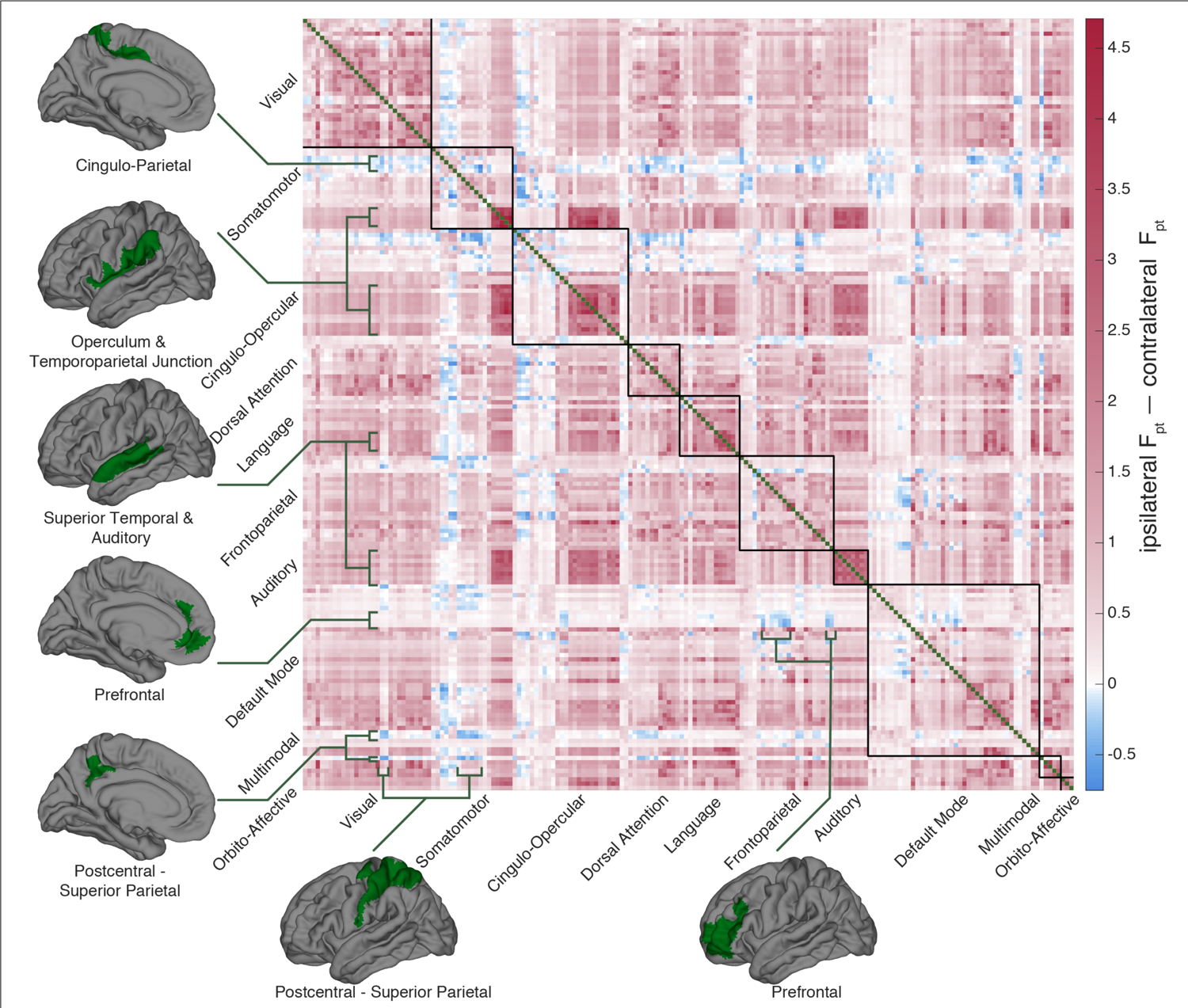
Interhemispheric connectivity. Differential connectivity between ipsilateral and contralateral connectivity. Greater Ipsilateral connectivity dominates and is indicated in red. Parcel-pairs with greater *contra*lateral connectivity than ipsilateral are blue. The green cortical patches show anatomic extent of parcel groups of notable contrast.

### With the exception of some language areas, most parcels are disproportionately connected to their contralateral homologs

The two hemispheres of the cortex have a high degree of functional and anatomical symmetry. It follows then that most regions will have greater connectivity to their contralateral homologs than other contralateral areas, in order to coordinate their overlapping processing tasks. This is hinted at by the visibility of the 180^th^, (or half-) diagonal in **figure 1A**. To further quantify this effect, for all 180 parcels we compared the connectivity between interhemispheric homologs to the mean of all other callosal connectivity. Bonferroni corrected, empirical 95% confidence intervals were estimated via bootstrapping with 2000 iterations. As detailed in extended data **figure 6-1** and visualized in **Figure 6**, 147 parcels are hyperconnected to their contralateral homologs, 18 are hypoconnected, and 15 have homologous callosal connectivity not significantly different than their callosal mean connectivity. Interestingly, parcels that are not hyper-connected to their contralateral homologs are concentrated within and adjacent to the language network, consistent with the greater degree of lateralization in these areas.

**Figure 6.**
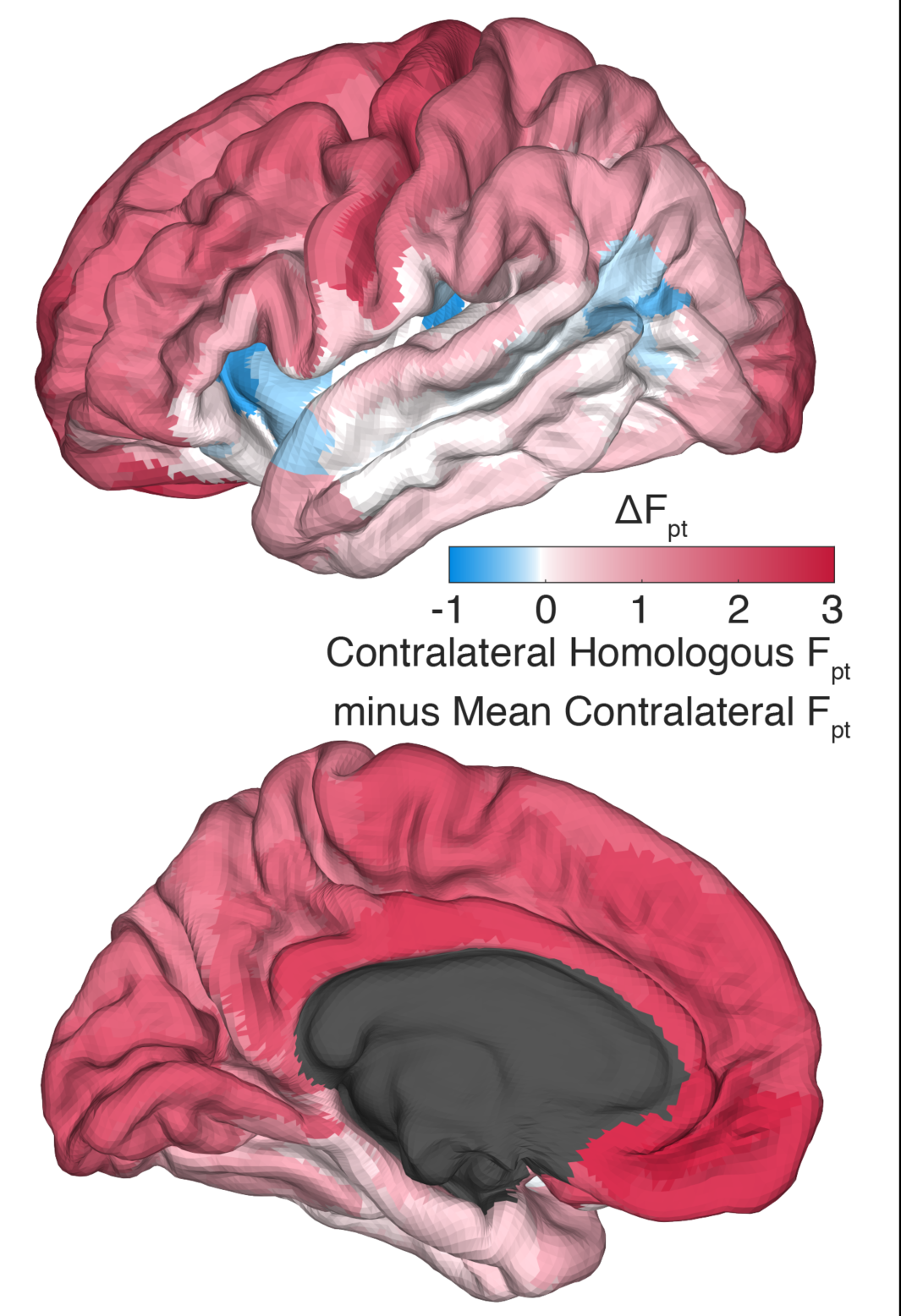
Contralateral homologs. Differential connectivity between contralateral homologous parcels vs the mean of all other contralateral parcels. Red indicates contralateral homologous connectivity greater than mean contralateral connectivity. Note that many language-implicated regions have relatively weak connectivity with their contralateral homologs.

**Figure 6-1.**
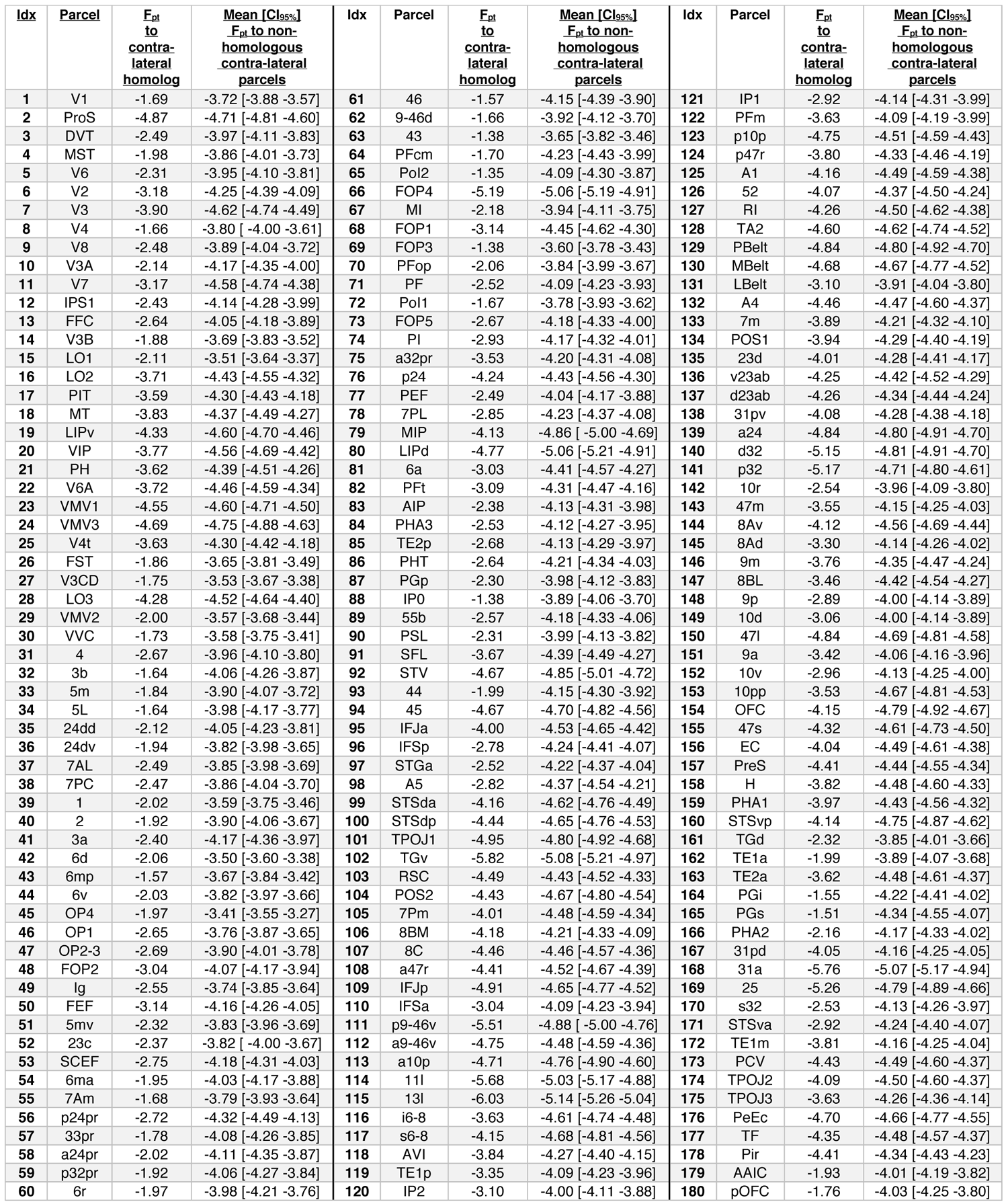
Differential connectivity between contralateral homologous parcels vs the mean of all other contralateral parcels. Confidence intervals are Bonferroni-corrected for multiple comparisons.

### The language network is hyper-connected at long distances and left lateralized

In order to investigate distance-resolved left laterality in connections among language-implicated cortex, pairwise connections were binned by fiber tract length in 15 mm increments. Within each bin, connections were grouped as being within the combined language and auditory network, or between the combined networks and the rest of the cortex. For each subject, the F_pt_ of grouped connections within each bin was averaged before being log-transformed. The grand-averages of these within-and between-language/auditory cortex in each distance bin for each hemisphere are shown in **figure 7A**. Bonferroni corrected, empirical 95% confidence intervals for these grand-averages were estimated via bootstrapping with 2000 iterations. Within-language connectivity is slightly attenuated at distances less that 100 mm, but strongly amplified at distances above 100 mm, especially ∼100-140 mm connections in the left hemisphere. A plurality of these are between frontal and temporoparietal language areas (18/45 connections between 100 and 140mm). The differential traces of between-vs. within-language connectivity (**Fig. 7B**) clearly show the left-hemisphere dominance of this effect.

**Figure 7.**
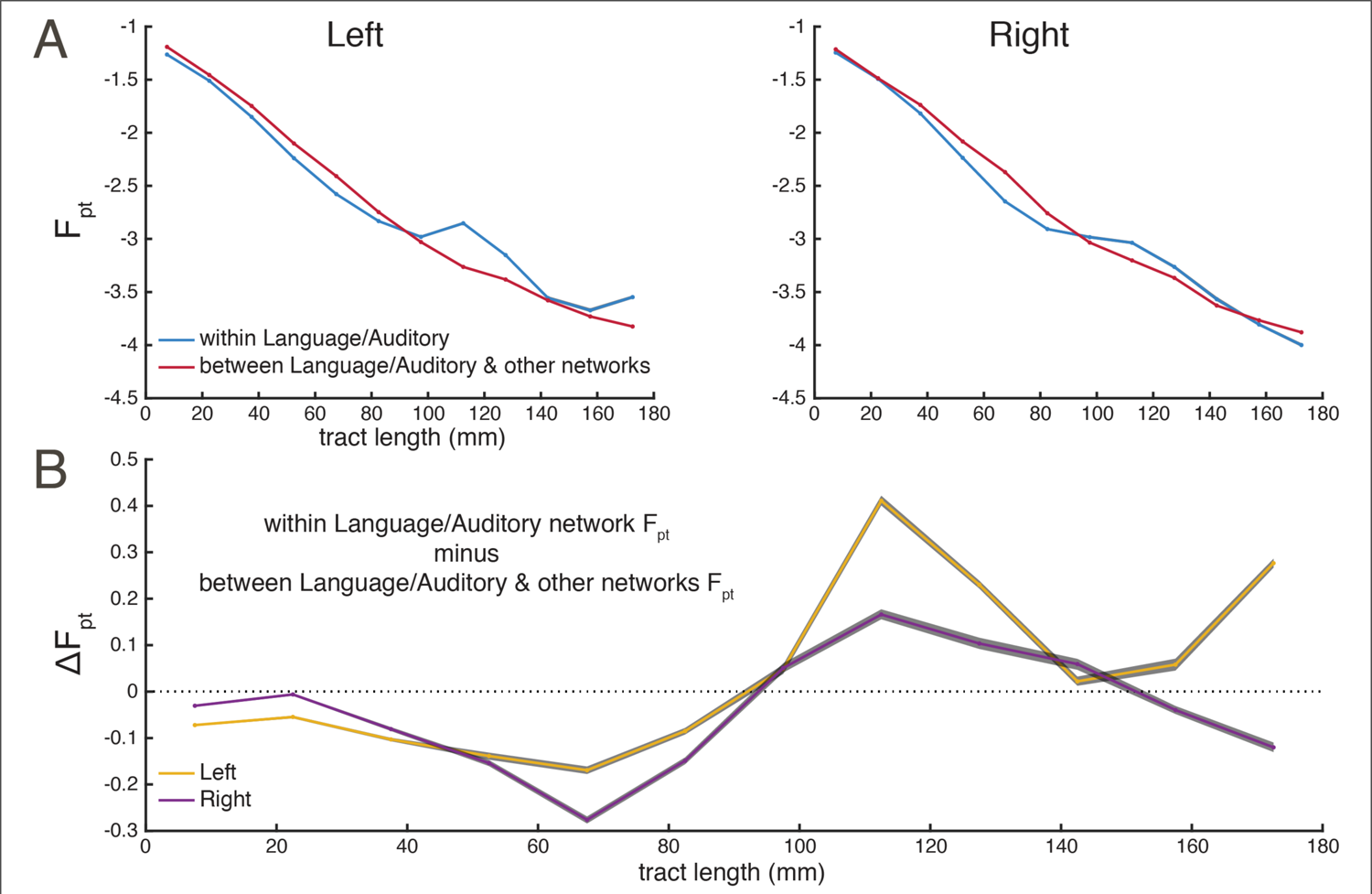
Language/Auditory network hyperconnectivity and left-lateralization. (**A**) Distance-binned connectivity within the language and auditory networks compared to connectivity between the language and auditory networks and other networks, separately for the left and right hemispheres (**B**) The differential trace for the within- and between-connectivity in both hemispheres. In both panels, gray patches show Bonferroni-corrected bootstrapped 95% confidence intervals across subjects.

### Connectivity is influenced by the cortical hierarchy

Hierarchy is a central organizing principle of the cortex (Burt et al., 2018; Felleman and Van Essen, 1991; Markov et al., 2014; Theodoni et al., 2020). Higher order areas, e.g. supporting abstract processing, have low myelination, and lower order areas, e.g. supporting unimodal sensory processing, have high myelination. Furthermore, areal myelination is indexed by the ratio between T1- and T2-wieghted MRI contrast (Glasser and Van Essen, 2011). The WU-Minn HCP 1200 release includes smoothed group-average myelination indices for all vertices in the 32k grayordinate template brain. These values were averaged for each parcel in the HCP-MMP1.0 atlas (Glasser et al., 2016) to yield a group-average parcel-wise index of myelination.

The relationship between cortical hierarchy and connectivity was assessed in two ways. We first examined whether regions of similar level in the cortical hierarchy are better connected, as predicted by (Barbas, 2015). An index of hierarchical similarity, F_|Δ myelination|_, was obtained for each pair of parcels by computing the pairwise difference in myelination between parcels and fractionally scaling it in the same manner as F_pt_, with smaller values indicating hierarchical closeness. The similarity matrix created by this derivation is shown in **figure 8-1**. Correlations were obtained for the left and right hemisphere as a whole as well as the colossal connections, **figure 8A**. In addition, for each of the twenty functional networks (10 per hemisphere) the Pearson correlation between the F_|Δ myelination|_ and F_pt_ for pairwise within-network connections was computed, see **figure 8B**. With the exception of the interhemispheric connections, calculations were performed on the hemispheres separately to avoid the collinearity introduced by hemispheric homology.

**Figure 8.**
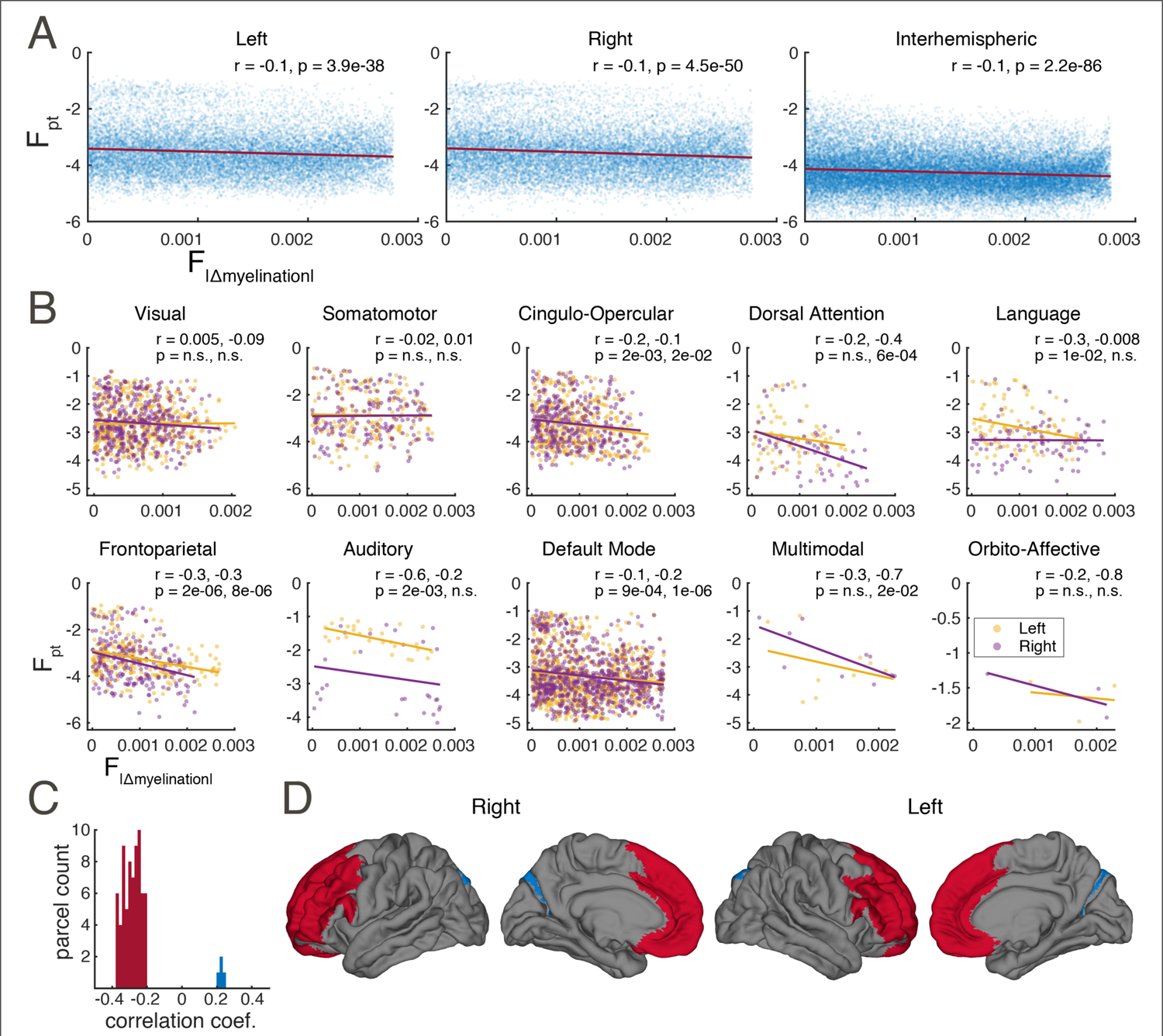
Connectivity is influenced by the cortical hierarchy. (**A**, **B**) Connectivity is strongly predicted by hierarchical similarity in some networks and modestly predicted overall. (**A**) All connectivity vs. myelination difference, including within- and across-network connections, for the left, right, and callosal connections. For both panels, each marker represents a parcel pair. (**B**) Within-network connectivity vs. myelination difference for 10 functional networks. Linear fits and correlation coefficients computed independently for the left and right hemisphere. A negative correlation indicates that parcels at similar hierarchical levels tend to be more connected. (**C**, **D**) Higher order prefrontal areas are better connected. (**C**) Histogram of correlation coefficients between areal myelination and F_pt_ connectivity to each parcel. Only significant coefficients after Bonferroni correction are shown. Most coefficients are negative indicating high connectivity to low-myelination (i.e., higher-order) areas. (**D**) Significant negative coefficients (red) map onto bilateral prefrontal cortex. Only the bilateral DVT and V6A are show positive significant correlations (blue).

**Figure 8-1.**
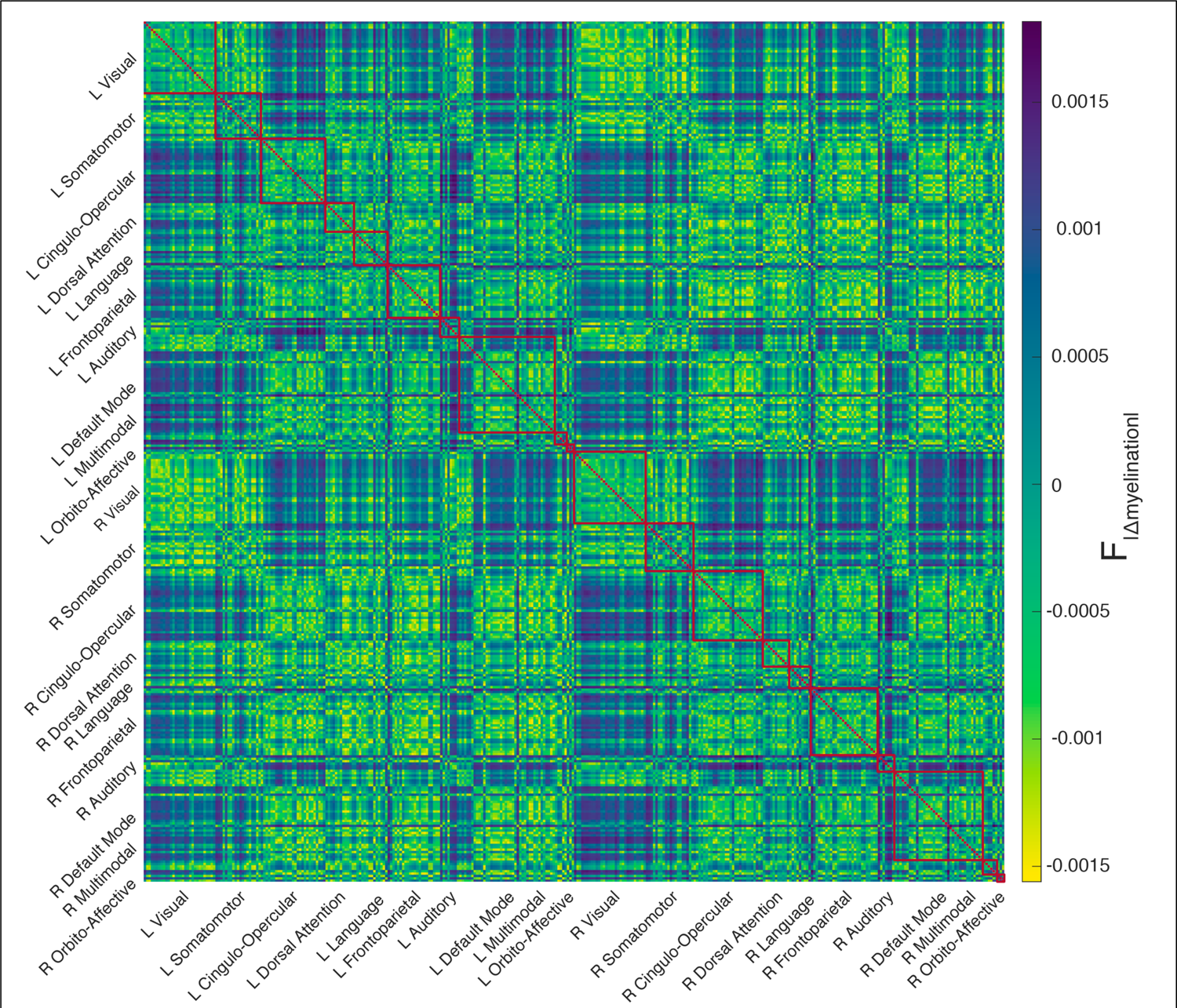
Myelination difference connectivity matrix. This provides an estimate for the difference in hierarchical level between cortical parcels. Values have been fractionally scaled. Note that the color scale has been reversed when compared to figure 1, as |Δmyelination| is inversely proportional to connectivity.

**Figure 8-2.**
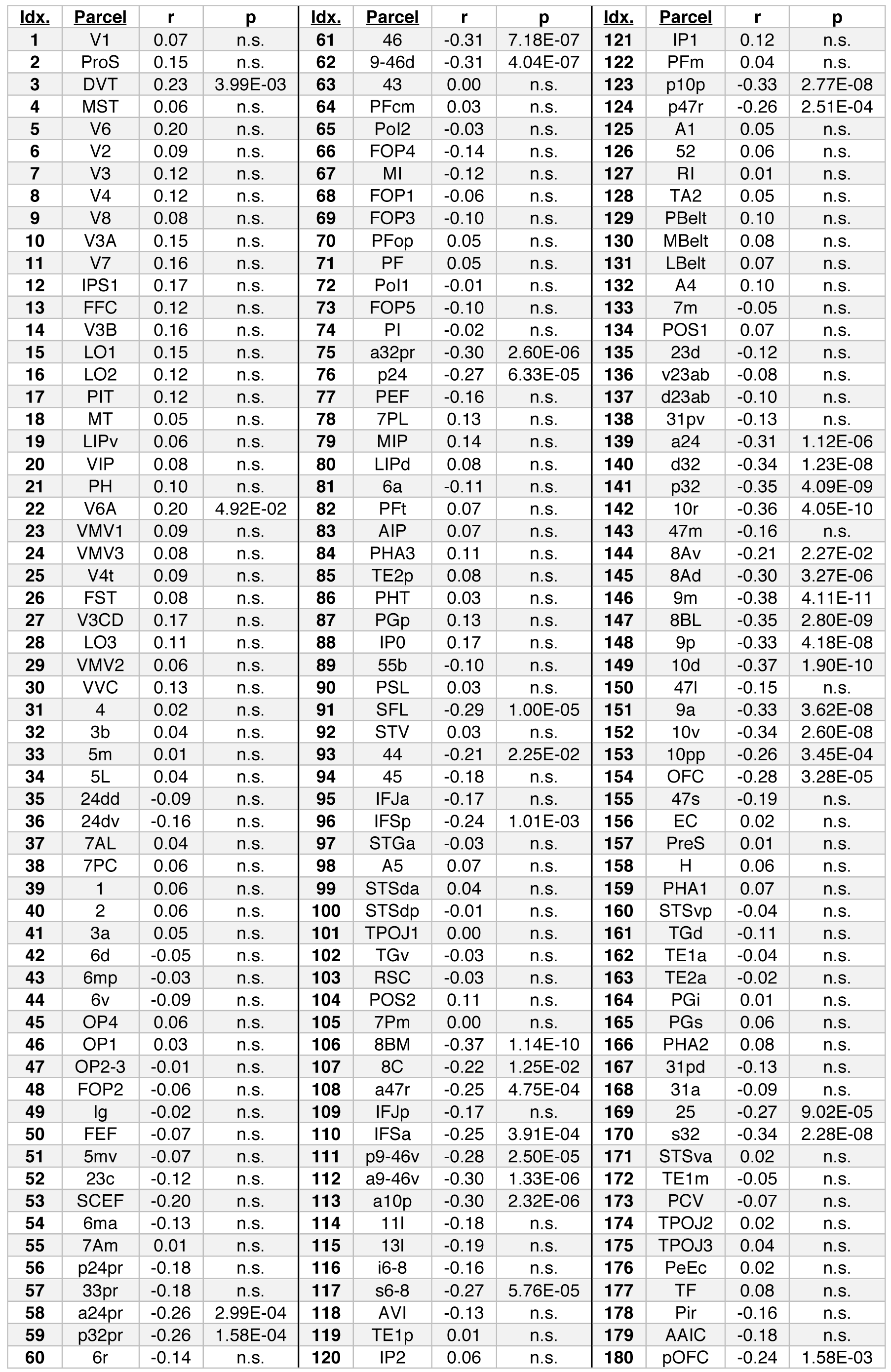
Pearson correlations between the F_pt_ from each left hemisphere parcel to all others and the target parcels’ myelination indices. p values are Bonferroni-corrected for multiple comparisons.

**Figure 8-3.**
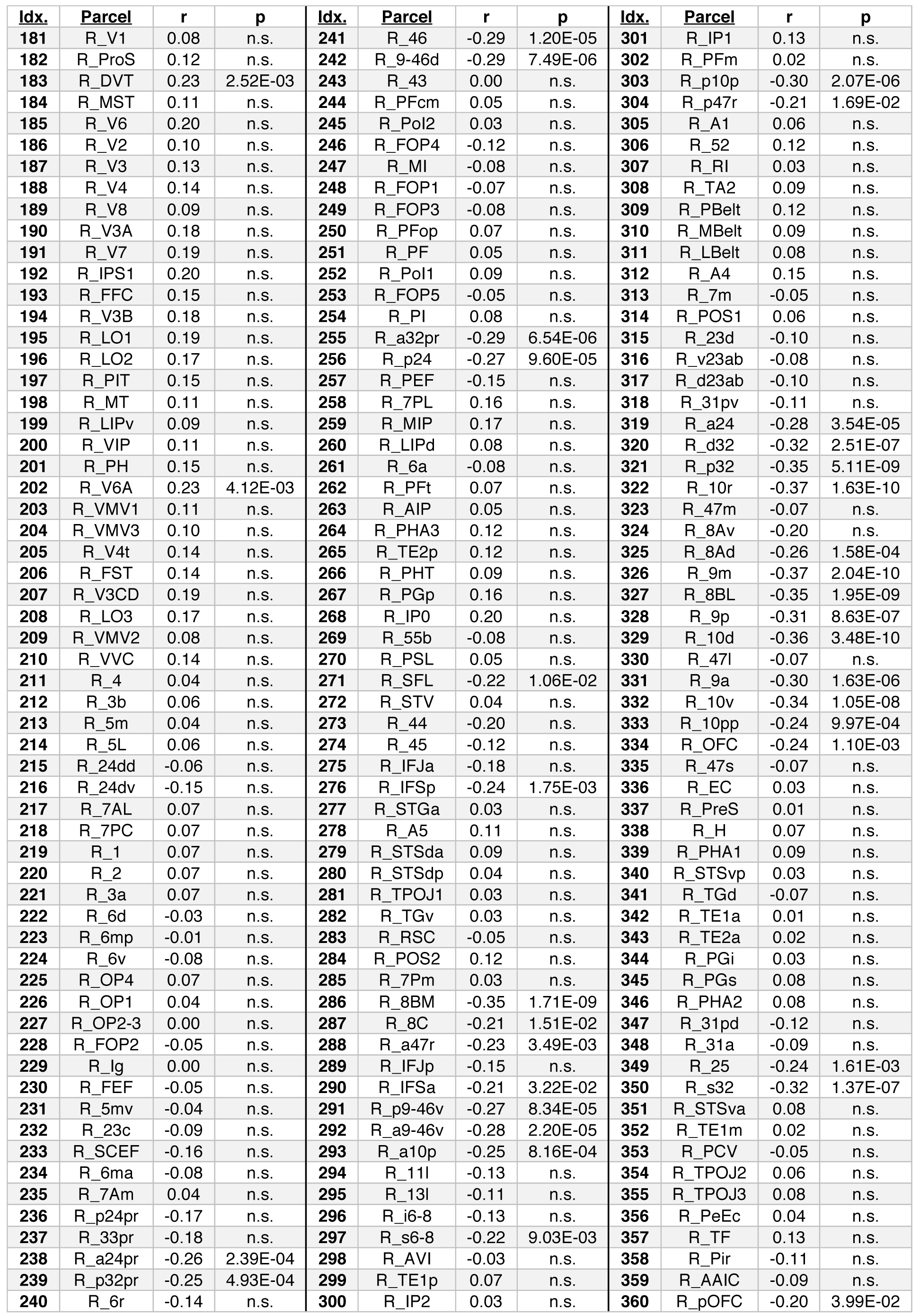
Pearson correlations between the F_pt_ from each right hemisphere parcel to all others and the target parcels’ myelination indices. p values are Bonferroni-corrected for multiple comparisons.

With the exceptions of the bilateral visual and somatomotor networks and right language network, for which there is convincingly no relationship, the preponderance of coefficients are negative, indicating that, on average, areas at similar levels of the cortical hierarchy are better connected. However, quantified in this way, the influence of hierarchy is modest, explaining about 1% of the variance in F_pt_ overall, though perhaps 10-30% in certain subsets of parcels, such as the left auditory and language networks. The left lateralization of the influence of hierarchy in these networks is striking, as is the right-lateralization of the dorsal attention network.

Secondly, we investigated whether a cortical region’s hierarchical level affected its overall connectivity. For each parcel, the Pearson correlation between the parcel’s F_pt_ to all other parcels and the parcel-wise index of myelination was computed. In other words, correlation between each row of the connectome matrix and the vector of myelination indices was obtained. After Bonferroni correction for multiple comparisons, 74 of 360 parcels (see extended data **figures 8-2, 8-3**) have connectivity significantly correlated to their myelination index and of these the vast majority (70) are negatively correlated, indicating that low myelination predicts high connectivity, see **figure 8C**. These areas form a contiguous bilateral prefrontal network as shown in **figure 8D**, indicating that prefrontal areas are more connected with higher cortical regions. The rare positively correlated exceptions are the left and right DVT and V6A.

### Probabilistic dMRI connectivity more closely resembles CCEPs than resting-state fMRI

In order to further contextualize the dMRI connectome, we compared it to existing connectivity matrices generated from two other brain mapping modalities: cortico-cortico evoked potential probability (CCEP) and resting-state fMRI correlation magnitude (rs-fMRI). As shown **figure 9A**, the qualitative pattern of rs-fMRI markedly differs from the other two modalities with proportionally stronger ipsilateral across-network connections and especially non-homologous contralateral connections, though the latter is somewhat obscured for CCEPs due to sparse spatial sampling. Over all connections, pairwise probabilistic dMRI connectivity values are nearly twice as linearly correlated to pairwise CCEP connectivity than to rs-fMRI connectivity (**fig. 9B**), and this contrast is equally evident in the ipsilateral connection within each hemisphere, see extended data **figure 9-1**. Contralateral connections were not examined in isolation as contralateral sampling for the CCEP modality is relatively rare.

**Figure 9.**
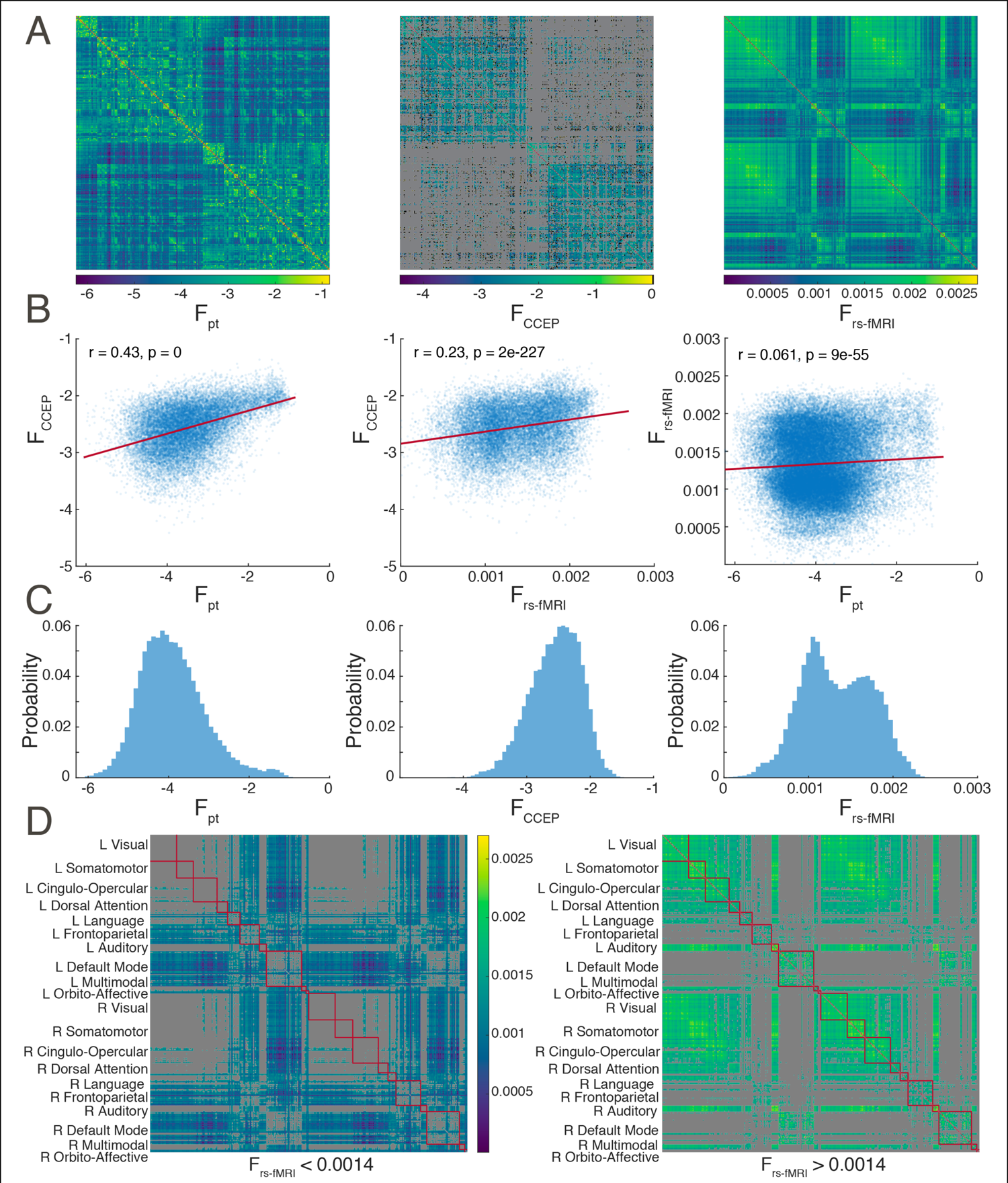
Probabilistic dMRI more closely resembles CCEPs than resting-state fMRI. (**A**) Connectivity matrices for probabilistic dMRI tractography, CCEP, and rs-fMRI. For CCEPs missing data has been colored grey and pre-log zero-strength connections black. (**B**) Correlations among the three modalities. The least-squares linear fit is shown in red. (**C**) Non-zero pairwise connection strength distributions. Note that rs-fMRI connectivity values, which are not log-transformed, display two modes, separated at 0.0014.(**D**) Cortical parcels displaying lower (left) and higher (right) modes of rs-fMRI connectivity.

**Figure 9-1.**
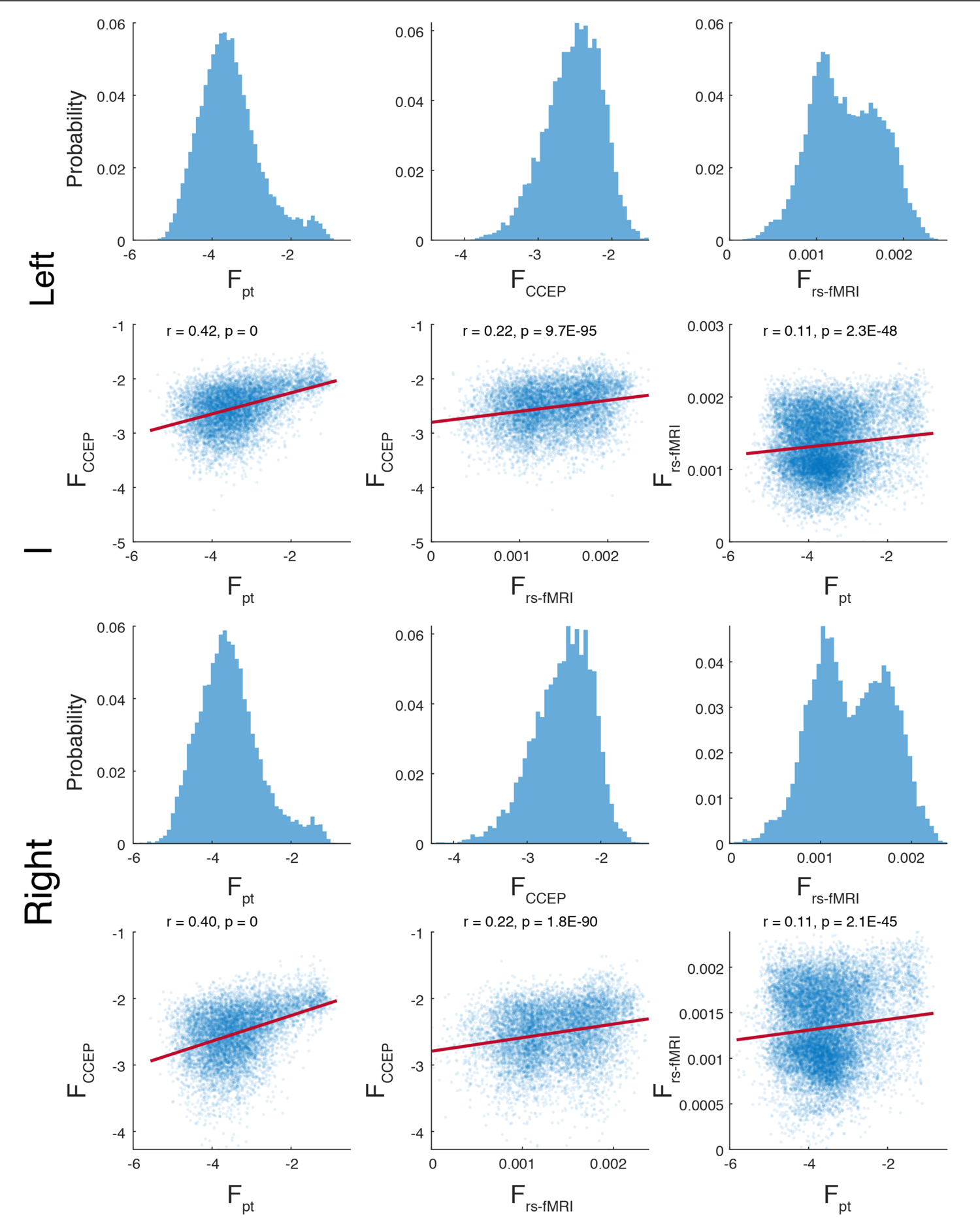
Within-hemisphere comparison of probabilistic dMRI tractography, CCEP, and rs-fMRI connectivity. For the left and right hemisphere, the distribution of pairwise non-zero connection strengths and correlations among the three modalities are shown. The least-squares linear fit is shown in red. All within-hemisphere findings are concordant with the overall findings, shown in **figure 9**.

When comparing the distributions of pairwise connectivity strength (**fig. 9C**), rs-fMRI again exhibits properties different than the other two modalities. While both dMRI and CCEP distributions skew in opposite directions (0.63 and −0.43, respectively), their strengths form unimodal log-normal distributions and thus shown with log-transformed values. In contrast, rs-fMRI connectivity values form a bimodal Gaussian-mixture distribution in linear space. The two modes were characterized by obtaining the maximum-likelihood fit (`fitgmdist`) of a 2-component Gaussian-mixture to the data, yielding a left mode (μ = 0.0011, σ = 8.1e-8) forming 63% of the distribution and a right mode (μ = 0.0017, σ = 8.1e-8) forming 37%, respectively. Splitting the rs-fMRI modes at the midpoint between their means (0.0014) and plotting their respective connectivity matrices (**fig. 9D**) reveals that the low-connectivity (left) mode consists primarily of connections between the default mode / frontoparietal networks and other regions of the cortex.

To further contrast the three connectivity modalities we computed six network theoretic metrics for each of the connectivity matrices: mean clustering coefficient (MCC), characteristic path length (CPL), global efficiency, gamma (normalized MCC), lambda (normalized CPL), small worldness, transitivity, and assortativity (see Appendix). Binarized network metrics were assessed after thresholding by edge weight (connectivity strength) at intervals of 0.1. Note that this lambda is unrelated to the exponential length constant reported above. To account for the order-based arbitrary treatment of equal edge weights when thresholding, the node (parcel) order was randomized 1000 times, and the mean metric values are shown. Empirical 95% confidence intervals for these means are too small to be shown at scale. Networks densities above 0.6 were not examined as the un-thresholded network density of CCEP connectivity matrix, treating missing data as non-connections, is less than 0.7. However, all measures appear to converge as binary network density approaches 1. As shown in figure 10, the MCC, CPL, global efficiency, small worldness, transitivity, and assortativity are markedly different for rs-fMRI connectivity than for CCEP and probabilistic dMRI tractography, whose metrics as a function of network density are more similar to each other. Normalizing by metrics computed for a random network with the same statistical makeup changes this pattern. For gamma the rs-fMRI and CCEP networks are more similar than either is to probabilistic dMRI tractography, and lambda rs-fMRI and probabilistic dMRI tractography are more similar than either is to the CCEP network. The high MCC, transitivity, and assortativity and low global efficiency of rs-fMRI relative to the other modalities may be indicative of strong, long-range correlativity beyond that predicted by anatomical connections.

**Figure 10.**
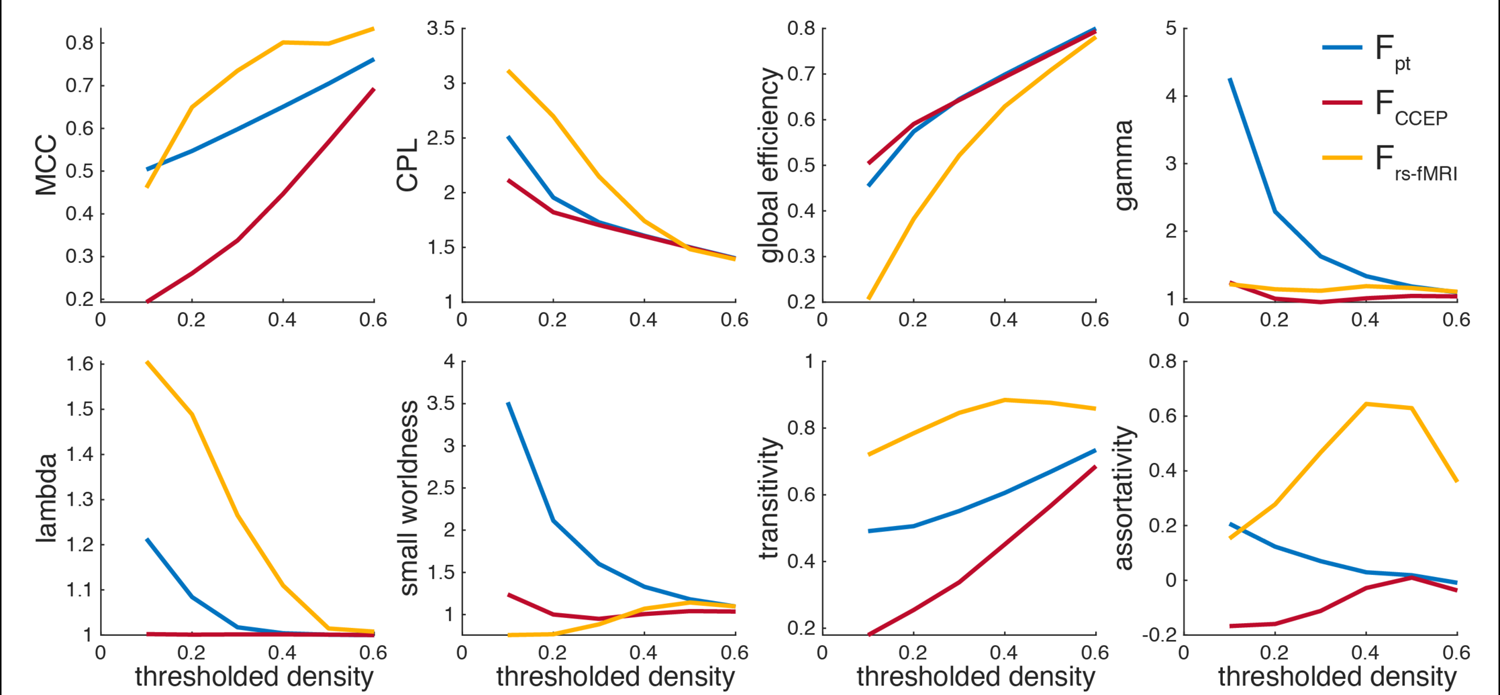
Network theoretic differences between the connectivity modalities. Binarized network metrics after thresholding by edge weight (connectivity strength).

## Discussion

In this study we compiled a whole-cortex structural connectome by applying probabilistic tractography to the diffusion MR volumes of 1065 subjects from the WU-Minn Human Connectome Project. We report a novel, complete, and high-dynamic-range connectivity matrix discretized into the 360 parcels of the HCP-MMP1.0 atlas and further arranged into 10 functional networks. It is shown that connectivity strength exponentially decays with fiber tract length, that the parts of the connectome with clear homology to macaques correspond reasonably to retrograde tracer mappings in that species, that contralateral homologs are hyperconnected, and that some connections within language-implicated cortex are stronger than expected and left-lateralized. While ipsilateral connectivity generally dominates, some regions have stronger contralateral connections. Inter-individual variability is relatively high for early visual cortex, whose connectivity co-varies across hemispheres. Cortical areas tend to be more connected with areas at similar levels of the cortical hierarchy, as indexed by their estimated myelination, particularly in prefrontal areas. Lastly, it is shown that probabilistic tractography connectivity more closely resembles that of CCEPs than rs-fMRI. In sum, we quantify a dMRI-based estimate of medium-to long-range anatomical cortico-cortical connectivity in a large normative sample.

Diffusion MR imaging and automated post-hoc tractography are powerful tools for the elucidation of cerebral connectivity. The defining advantages of these techniques are non-invasiveness and large field-of-view, enabling whole-brain mapping in humans. However, dMRI does have significant limitations when compared to histological fiber tracing, EM microscopy, or stimulation. The most obvious of these is insensitivity to whether underlying axons are anterograde or retrograde, as evidenced by the symmetry of the connectivity matrix. The anisotropic diffusion of water molecules occurs in both anterograde and retrograde directions. Thus, the true one-way connectivity between two areas could be anywhere between none to all of the symmetric diffusion connectivity. Another important limitation is spatial resolution. While the 1.25 mm isotropic voxels achieved by the WU-Minn dMRI protocol are smaller than those of most studies (Jeurissen et al., 2019), they are still more than three orders-of-magnitude larger than the typical submicron axon diameter (Liewald et al., 2014; von Keyserlingk Graf and Schramm, 1984). This discrepancy is particularly impactful when fiber orientations are not consistent within a voxel, i.e. crossing fibers. Probabilistic diffusion tractography (Behrens et al., 2007) partially ameliorates the issue by modeling the probability distribution of orientations and accounting for uncertainty, but ultimately dMRI with current technology is a meso- to macroscale technique. Direct histological validation of dMRI techniques is uncommon, but has been performed for probabilistic tractography *in vitro* in pigs (Dyrby et al., 2007) and macaques (Donahue et al., 2016; Jbabdi et al., 2013), with the latter two studies using the same probtrackX algorithm as the current study (Behrens et al., 2007). We have extended these validations with a between-species comparison (**Fig. 4**).

Of the several families of dMRI tractography algorithms available, we selected local, probabilistic tractography (Behrens et al., 2007). The WU-Minn HCP makes available the bedpostX precursor files and creating a probabilistic tractography connectome was always a stated component of the WU-Minn HCP project (Van Essen et al., 2013; Van Essen and Ugurbil, 2017). That such a connectome has not yet been released for these data may be due to the immense computational challenge of performing these analyses at the scale of the HCP. An advantage of probabilistic tractography is its sensitivity to minor, or low-probability connections. Deterministic dMRI tractography connectomes typically have low network densities, e.g. 0.18 (Mori et al., 2008) or 0.23 (Cui et al., 2019), when compared to histological fiber tracing in macaques, 0.66 (Markov et al., 2014), and this is likely a lower bound as such tracing is subject to false-negatives due to imperfect dye uptake and incomplete cortical sampling. This suggests the deterministic dMRI connectomes are missing weaker connections. On the other hand, dMRI in general and probabilistic tractography in particular has been found vulnerable to false-positive connections (Maier-Hein et al., 2017). This exchange of specificity for sensitivity (Sarwar et al., 2019; Zalesky et al., 2016) is consistent with our very high group-average network density of 1.0 and the likely presence false-positive connections, and is thus an important caveat to the data presented here. In cases where false-negative connections are less concerning than false-positive connections, such as topological analyses (Zalesky et al., 2016), subsequent users of these data may opt to threshold the connectivity matrix by either connection strength or consistency (Roberts et al., 2017), see **figure 3**.

When constructing this connectome, we divided the cortex into 180 parcels per hemisphere following the HCP-MMP1.0 atlas (Glasser et al., 2016). To ease interpretation, we further organized the parcels into 10 functional networks modified from (Ji et al., 2019). These networks were created by applying iterative Louvain clustering (Blondel et al., 2008; Rubinov and Sporns, 2010) and other criteria to HCP resting state fMRI data. While these fMRI-defined network definitions correspond reasonably to the structural connections reported here, there are exceptions. The operculum and temporoparietal junction, in particular, appears to be a structurally distinct area that has been folded into several functional networks (Ji et al., 2019). However, this contiguous region forms the lateral salience network in (Barnett et al., 2020) which similarly applied a very similar methodology to a non-HCP cohort. Like many cortically-focused studies, we used a surface-based methodology to define these areas, with seed and target regions constrained to the white-matter – gray-matter interface. This approach reduces the overrepresentation of major bundles (Jeurissen et al., 2019), enables the automated assessment based on inter-subject homology (Fischl et al., 1999), facilitates comparison to other cortical datasets, and is true to the anatomical nature of the cortical ribbon.

Unfortunately, the subcortex and cerebellum are omitted in this analysis, as are short-range, often unmyelinated, intra-parcel connections. While the inclusion of the thalamic radiations, in particular, is a merited future extension of this connectome, the small size of subcortical structures relative to diffusion imaging voxels, the nuclear (as opposed the sheet-like) organization of subcortical structures, and complex geometry of the subcortical white matter — gray matter interface (e.g. the internal medullary lamina of the thalamus), all render the challenges and methods for obtaining subcortical tractography substantially distinct from those of cortico-cortico tractography.

The HCP-MMP1.0 atlas used was selected because of its wide adoption, symmetry, and high parcel count. Furthermore, the parcels are based on multiple functional and anatomical criteria and are consistent with previous functional parcellations in human and non-human primates (Felleman and Van Essen, 1991; Glasser et al., 2016). Because the parcels are relatively small and informed by function, erroneous averaging of disparate connections, a connectomic extension of the partial volume artifact, is minimized. However, this comes at the cost of non-uniformity in both parcel area and shape. Methodologically, parcels are assembled from vertices on the tessellated cortical surface. A future vertex- or voxel-based connectome, while computationally challenging, would have the distinct advantage of being readily reformulated into any arbitrary surface-based parcellation scheme.

We found that pairwise connectivity between cortical parcels exhibits an exponential decay rule with respect to fiber tract distance with a length constant λ of ∼23 mm (∼33 mm for callosal connections). While a tight exponential relationship between probabilistic diffusion tractography strength and fiber length has been previously reported, (Roberts et al., 2016), this study did not report the observed λ or release its data. Histological studies in non-human primates (Donahue et al., 2016; Markov et al., 2013; Theodoni et al., 2020) consistently show exponential connectivity decay with distance. Such a rule when combined with a roughly Gaussian distribution of interareal distances explains the observed log-normal distribution of connectivity strength (Markov et al., 2013). Histological data indicate a λ of about 3.33 mm for marmosets (Theodoni et al., 2020) and 5.55 mm for macaques (Markov et al., 2013). Across species, there appears to be a linear relationship between the logs of λ and total gray matter volume, predicting a human λ of 10 mm (Theodoni et al., 2020). While methodological differences between diffusion and histological tractography cannot be completely ruled out, Donahue and colleagues found similar λ for the two methods in macaques (Donahue et al., 2016). Our results suggest that, compared to other species, human cortical areas are exceptionally well connected relative to their cortical volume, reflected in a disproportionately long λ. Conservatively restricting the exponential fit to only the most consistent quintile of connections (**Fig. 3D**) yields a λ of ∼28 mm, further accentuating the proportional long-range hyperconnectivity of humans.

Geometric scaling strongly constrains cortico-cortical connectivity in humans. Considering primate brains increasing in diameter d, volume and number of cortical neurons increases by d^3^,(Ventura-Antunes et al., 2013), so arriving at a constant probability of connection between any two neurons would require d^6^ axons, and since they would need to be about d times as long, this would require a volume proportional to d^7^, or more if axonal diameter is increased to maintain a relatively constant latency of communication (Wang et al., 2008). However, the actual white matter volume is less than d^4^ (Zhang and Sejnowski, 2000), and consequently the probability of cortico-cortical connectivity must be highly limited in humans. The relatively long λ in humans we report reduces even further the number of connections which can be accommodated within the available white matter volume. A consequence of fewer but longer connections would be reduced metabolic cost, inasmuch the cost of an action potential is 1/3 axonal transmission (proportional to length) and 2/3 synaptic transmission (Lennie, 2003). The low firing rate of human pyramidal cells (Chan et al., 2014) would also reduce the metabolic cost of their axons. These observations are consistent with the proposal that the metabolic costs of cortico-cortical connections may help constrain their organization in the primate brain (Ercsey-Ravasz et al., 2013). Given this strong correlation of connection strength with distance, as well as the bias of tract-tracing techniques toward shorter, less geometrically complex connections (Jeurissen et al., 2019), there may be some merit in regressing out the effect of tract length when evaluating the relative connectivity of different cortical areas. However, the considerations enumerated above imply a strong evolutionary selection to place cortical parcels which require high connectivity to perform their calculations to be situated in direct physical proximity to each other. The patterns of relatively long distance connectivity identified here thus must be viewed as minor deviations from an overall strong tendency favoring local connectivity, a conceptualization consistent with the view of the cortex as a spatially embedded small world network.

One striking deviation from the distance-based connectivity was the left-lateralized hyper-connectivity between language areas, and specifically between posterior and anterior language areas. This connectivity presumably passes, completely or in part, through the classical language pathways (reviewed in (Dick and Tremblay, 2012)). The lateralization we observed may then reflect that of the arcuate and inferior longitudinal fasciculi which connect the same structures and show significant left lateralization in humans but not macaques (Eichert et al., 2019; Panesar et al., 2018). Left-lateralization of the arcuate fasciculus develops late (Lebel and Beaulieu, 2011), and is sensitive to the presence, quality and quantity of early language experience (Cheng et al., 2019; Romeo et al., 2018). More generally, many of the connectivity patterns observed here could be the indirect result of co-activation of the connected parcels (Mount and Monje, 2017). The left-lateralized ipsilateral connectivity may be compensated by a relative lack of callosal connections from the same areas, under the hypothesis that the total connectivity is constrained.

A more general factor that might induce deviations from a distance-based connectivity rule may be the principle of hierarchical organization. It has been proposed that distant areas with similar laminar properties, and thus of similar hierarchical order may have privileged connections (Barbas, 2015). Across the entire cortex we find that myelination similarity explains a significant but small amount of the overall variance. However, there are regions where the influence of hierarchical position is more pronounced including the right dorsal attention and left auditory/language networks. The observed hyperconnectivity and high degree of lateralization in these regions may be a consequence of the low-latencies necessary for the functions they underly. More broadly, the effects of transmission latency constraints on neuroanatomy and conduction delay on large-scale physiological recordings are an emerging area of study in human neuroscience (Muller et al., 2018). Latency is a hybrid structural– functional property of connectivity, and might in future be quantified using the latency of cortico-cortical evoked potentials (CCEP).

By emphasizing the cortical connectivity matrix over the white matter bundles *per se* and organizing the matrix into the widely adopted HCP-MMP1.0 atlas (Glasser et al., 2016), the structural connectome reported here enables ready comparison to other structural, functional, and hybrid connectomes. As an example, we compared the probabilistic tractography connectivity to exist resting-state fMRI (rs-fMRI) (Van Essen et al., 2013) and CCEP (Trebaul et al., 2018) connectivity matrices and found that our dMRI-inferred structural connectivity better reflects CCEP probability than rs-fMRI connectivity in both linear and network-theoretic comparisons, despite the dMRI and rs-fMRI cohorts being highly overlapping. This is not unreasonable, as functional correlations are to varying degrees neurobehavioral state-dependent and far more spatiotemporally dynamic than structural connections. Furthermore, although resting-state functional connectivity is constrained by anatomical networks and can be partially predicted by them (Honey et al., 2009), indirect connections or parallel processing of stimuli in different areas can produce correlated activity even in the absence of direct anatomical connections. One notable example of the latter may be inter-hemispheric connectivity. While we did find hyperconnectivity between inter-hemispheric homologs when compared to other callosal connections, anatomical interhemispheric connectivity on the whole is much weaker than found in rs-fMRI. CCEPs, being directed by clinical requirements, have poor inter-hemispheric sampling, but we found that even among ipsilateral connections, rs-fMRI is still less similar to CCEP than probabilistic tractography. These inter-modal connectivity comparisons are not intended to be comprehensive. The HCP cohort also includes source-localized resting-state magneto-encephalography (MEG) (Larson-Prior et al., 2013), which could be used to examine the degree to which the functional connectivity of various frequency bands corresponds to anatomical connectivity. Furthermore, neuropsychological metrics, including the NIH toolbox (Gershon et al., 2013), and genotypic data (<u>dbGaP phs001364.v1.p1</u>) are also available for this cohort, enabling future studies of the interplay between cortical connectivity, cognition, and genetics.

The Human Connectome Project was a scientific undertaking of visionary scope and ambition. Its commitment to open science and accessibility of data by the public enabled this study and will continue to facilitate further studies for years to come. Emerging clinical applications of brain connectomics will be underpinned by a strong base of normative data for comparison. The whole-cortex probabilistic diffusion tractography connectome reported here fulfills a key goal outlined in the project’s conception and we hope it will empower yet further study of the myriad and beautiful web of connectivity that the human brain embodies.

### Data availability

Individual and group average connectivity matrices as well as all other figure source data can be found at https://doi.org/10.5281/zenodo.4060485 (https://zenodo.org/record/4060485). The preprocessed HCP data using in this study was retrieved from https://db.humanconnectome.org and the preprocessing code used to create these files is available at https://github.com/Washington-University/HCPpipelines. The source code for FSL, including probtrackx2 is available from https://fsl.fmrib.ox.ac.uk/fsl/fslwiki/FSL. Network theory measures were computed with the brain connectivity Matlab toolbox whose source code is available from http://www.brain-connectivity-toolbox.net.

